# Plasmid-encoded host reprogramming promotes plasmid dissemination

**DOI:** 10.64898/2026.07.07.736705

**Authors:** Chloé Virolle, Sibylle Ferrarin, Gaël Panis, Yannick Baffert, Annick Dedieu-Berne, Jérémy Guérin, Julien Cayron, Daouda A K Traoré, Sofía Martínez-Absalón, Rania Zenati, Frédéric Delolme, Adeline Page, Sarah Bigot, Yoshiharu Yamaichi, Allison Lopatkin, Patrick H. Viollier, David Burstein, Laurent Terradot, Christian Lesterlin

## Abstract

Conjugative plasmids are major drivers of antibiotic resistance dissemination, yet how newly transferred plasmids establish in recipient cells remains poorly understood. Here we investigate YfjB, a previously uncharacterized conserved leading-region protein, which is zygotically induced immediately after plasmid entry and acts specifically during the earliest post-transfer stages. Multi-omics analyses reveal that YfjB reprograms host transcription, triggering extensive metabolic rewiring that compensates the transient fitness cost of plasmid acquisition. Structural analyses show that YfjB is a ParB-like protein containing a CTP-binding domain and a helix-turn-helix DNA-binding motif, linked to a previously uncharacterized dimerization module that forms a V-shaped clamp-like architecture compatible with DNA loading. Consistently, live-cell imaging reveals nucleoid-associated foci in transconjugants, and ChIP-seq identifies multiple chromosomal binding sites. We therefore rename the protein HerB (Host Expression Reprogrammer, ParB-like). More broadly, our findings reveal how mobile genetic elements facilitate their dissemination by transiently subverting host physiology.

## Introduction

Conjugative plasmids are major drivers of antibiotic resistance dissemination, yet how newly transferred plasmids establish in recipient cells remains poorly understood. While the molecular machinery that mediates plasmid processing and transfer in donor cells has been extensively characterized, the molecular events that occur in recipient cells immediately after plasmid entry remain far less understood. This early phase, during which the incoming plasmid must establish itself in a naïve host, represents a critical bottleneck for successful horizontal transfer. Understanding this process may therefore illuminate the plasmid-host dialogue that shapes host range and dissemination trajectories, with broad clinical and biotechnological implications.

Following conjugative transfer, plasmid DNA enters recipient cells as a single-stranded molecule that must rapidly evade host defence systems and engage host replication and transcription machinery to ensure stabilization and inheritance. Increasing evidence indicates that these early challenges are addressed by functions encoded within the leading region, the first plasmid segment transferred into the recipient cell during conjugation. Leading-region genes are typically expressed immediately after plasmid entry through a process known as zygotic induction, driven by promoters that are active on single-stranded DNA and silenced once the plasmid is converted into double-stranded form^1–3^. Comparative analyses have shown that leading regions are enriched in functions that protect incoming plasmids from host defence mechanisms, including inhibitors of restriction-modification systems, anti-CRISPR proteins, suppressors of the SOS response and protective DNA methyltransferases^4–7^. These observations suggest that leading regions constitute specialized genetic modules dedicated to facilitating the earliest stages of plasmid establishment. However, despite their widespread conservation across conjugative plasmids, the functions of many leading-region genes remain unknown.

The F plasmid is both a classical model of bacterial conjugation and the prototype of widespread F-like plasmids that drive the dissemination of antibiotic resistance among Enterobacterales ^8–10^. Within its leading region lies *yfjB*, a gene that is zygotically induced immediately after plasmid entry and predicted to encode a ParB-like protein of unknown function. Canonical ParB proteins mediate faithful chromosome and low-copy plasmid partitioning through interaction with the ATPase ParA and binding to centromere-like *parS* sites. Although most characterized ParB proteins function in replicon segregation, several non-canonical ParB-family members have been evolutionarily repurposed for diverse cellular processes, including nucleoid occlusion and replication control (Noc)^11,12^ chromosome organization (PadC)^13^, regulation of integrative and conjugative element transfer competence (BisD)^14^, and fertility inhibition by conjugative plasmids (Osa-like proteins)^15^. Here, we identify a previously unrecognized function for a ParB-like protein. Using live-cell imaging, structural biology and multi-omics approaches, we demonstrate that YfjB is produced immediately upon plasmid entry into recipient cells, associates with the host chromosome and triggers extensive transcriptional and metabolic reprogramming. This transient host reprogramming primes recipient cell physiology for plasmid establishment, thereby alleviating the transient fitness burden of plasmid acquisition and promoting plasmid dissemination. Given this activity, we hereafter refer to YfjB as HerB (Host Expression Reprogrammer, ParB-like), a leading-region factor that transiently reprograms host physiology during the earliest stages of plasmid establishment.

## Results

### HerB forms nucleoid-associated foci in newly formed transconjugants

We previously demonstrated that HerB is zygotically induced immediately upon plasmid entry into recipient cells^1,16^. Using the same live-cell fluorescence microscopy system, we examined the accumulation and intracellular localization of HerB in newly formed transconjugants. In this system, donor cells carried an F *herB-sfgfp* plasmid encoding a C-terminal superfolder GFP fusion and a *parS* site positioned adjacent to the origin of transfer (*oriT*) (Fig. 1a). Recipient cells produced cytoplasmic mCherry-ParB, which remained diffusely distributed until plasmid acquisition. Upon plasmid transfer and conversion to dsDNA, mCherry-ParB was recruited to the *parS* site, producing a red focus that marks transconjugants (Fig. 1b).

**Figure 1.**
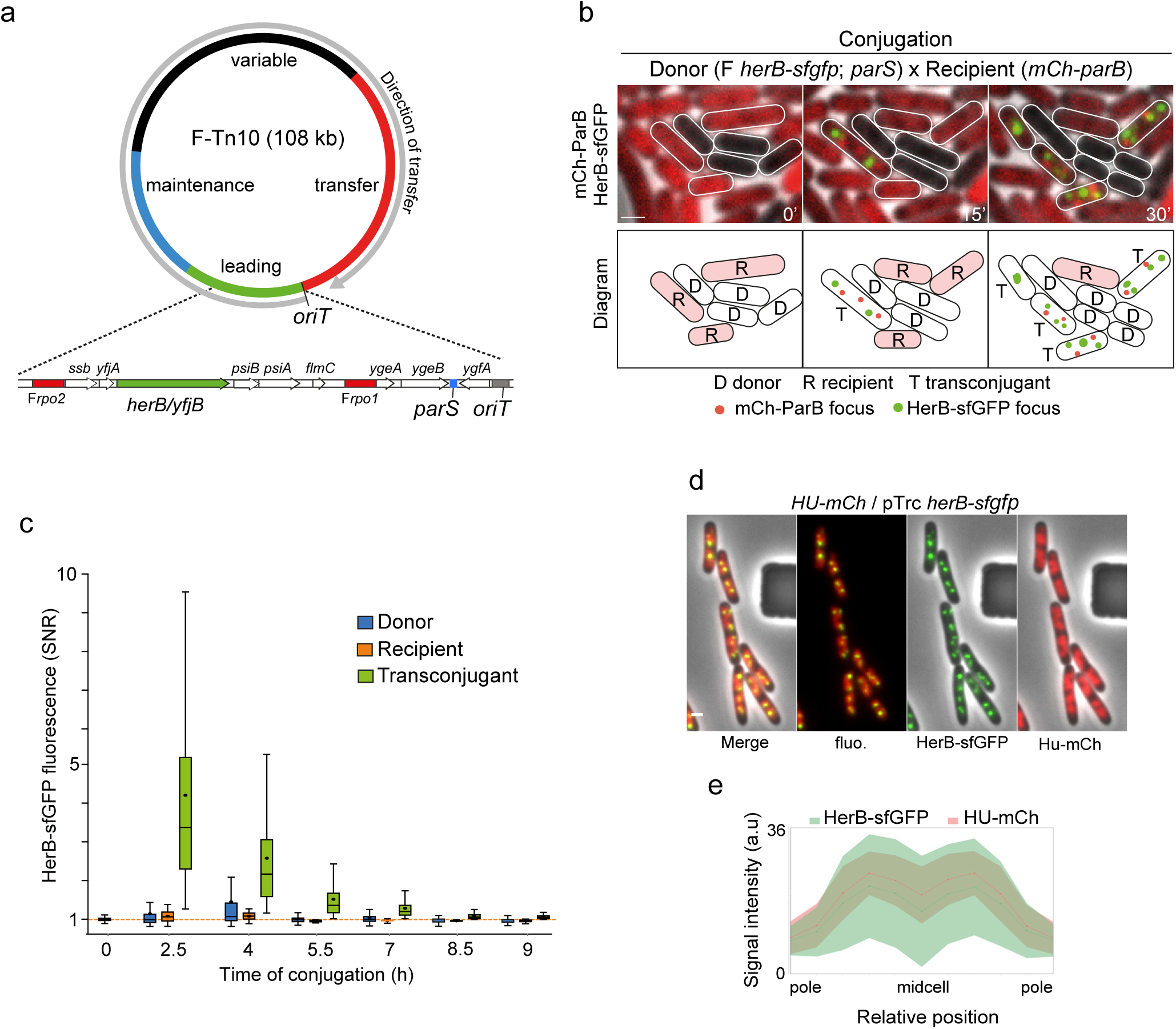
Transient production and intracellular localisation of HerB in transconjugant cells. **(a)** Genetic map of the F-Tn*10* plasmid showing the origin of transfer (*oriT*) and the direction of transfer (grey arrow), as well as the leading (green), maintenance (blue), variable (black) and *tra* (red) regions. The map of the leading region is shown in greater details below, with the *herb/yfjB* gene in green, the F*rpo1* and F*rpo2* single-stranded promoters in red, and the *parS* insertion site in blue. **(b)** Time-lapse microscopy images of conjugation between donor cells (D) carrying the F-Tn*10 parS, herB-sfgfp* plasmid and recipient cells (R) producing mCh-ParB. Transconjugant cells (T) that have acquired the plasmid exhibit red mCh-ParB/*parS* foci and HerB-sfGFP foci. A corresponding schematic is shown below. Scale bar, 1 μm. **(c)** Box plot of HerB-sfGFP intracellular fluorescence in donor, recipient and transconjugant cells over nine hours of conjugation. The median and interquartile range (Q1-Q3) are indicated by the box, the minima and maxima by the whiskers, and the mean by a black dot. The orange dashed line indicates the mean background fluorescence measured in recipient cells before mixing with donor cells. **(d)** Microscopy images of the cells producing HU-mCh from the chromosome and HerB-sfGFP from the pTrc *herB-sfgfp* plasmid within a microfluidic chamber. Scale bar, 1 μm. (**e**) Quantitative analysis of HU-mCh and HerB-sfGFP fluorescence distributions along the normalised long axis of cells grown in M9-CAA glucose medium. Lines represent the mean, and the coloured area represent the standard deviation (s.d.).

Quantitative analysis showed that HerB-sfGFP fluorescence was undetectable in donor cells, consistent with control of *herB* by F*rpo2* single-stranded promoter that is inactive when the plasmid is maintained in double-stranded form (Fig. 1b,c). In contrast, HerB-sfGFP formed bright, dynamic intracellular foci specifically in transconjugants (Fig. 1b; Extended Data Fig. 1a, Supplementary Video S1). These foci were positioned near quarter-cell locations and increased in number with cell length (1-5 per cell; mean 1.6 ± 0.7), indicating that HerB focus formation is coordinated with cell-cycle progression (Extended Data Fig. 1b-d). HerB-sfGFP fluorescence remained detectable for approximately seven hours, indicating that the protein persists for several generations before being diluted or degraded with in the transconjugant lineage (Fig. 1c).

Because HerB production from the F plasmid occurs only in transconjugants within mating mixtures, limiting population-level analyses, we constructed a pTrc99a *herB-sfgfp* plasmid enabling ectopic expression throughout the cell population. Quantitative snapshot microscopy showed that HerB-sfGFP expressed from this construct in cells lacking the F plasmid accumulated to levels comparable to those observed in F *herB*-*sfgfp* transconjugants (Extended Data Fig. 1e), and displayed a similar localization pattern (compare Extended Data Fig. 1a,b with 1f,g). Importantly, deletion of *herB* from the F plasmid did not alter its intracellular positioning (Extended Data Fig. 1h-k). Together, these observations indicate that HerB intracellular localization is independent of the presence of the F plasmid, and conversely that plasmid positioning does not depend on HerB.

To further characterize HerB localization, we examined cells expressing the nucleoid-associated protein HU-mCherry^17^. HerB-sfGFP fluorescence overlapped with HU-mCherry, indicating that HerB molecules are confined to the nucleoid region (Fig. 1d,e). This nucleoid-associated localization was maintained throughout the cell cycle and across growth conditions (Supplementary Video S2, Extended Data Fig. 1l). Treatment with rifampicin, which induces transcription arrest and nucleoid decompaction, caused a rapid disappearance of HerB foci. Following rifampicin washout, restoration of normal nucleoid organization was accompanied by the reappearance of HerB foci (Supplementary Video S3). Moreover, HerB foci were absent from DNA-free cells generated by complete chromosome degradation (Extended Data Fig. 1m)^18,19^. Together, these observations indicate that HerB localization depends on higher-order nucleoid organization rather than stable intracellular anchoring sites or the presence of the F plasmid, consistent with direct association with chromosomal DNA.

### Structural architecture of the ParB-like protein HerB

HerB is frequently annotated as a ParB-like protein (ParB2), as residues 36-243 display strong homology to members of the ParB/RepB/Spo0J family (Fig. 2a) (BLASTP E-value 6.07 × 10⁻²⁷). ParB typically creates homodimers and feature a N-terminal domain (NTD) that bind and hydrolyse CTP, a DNA binding domain (DBD) containing a Helix-Turn-Helix (HTH) motif and a C-terminal domain (CTD) mediating dimerization and aspecific DNA binding.

**Figure 2.**
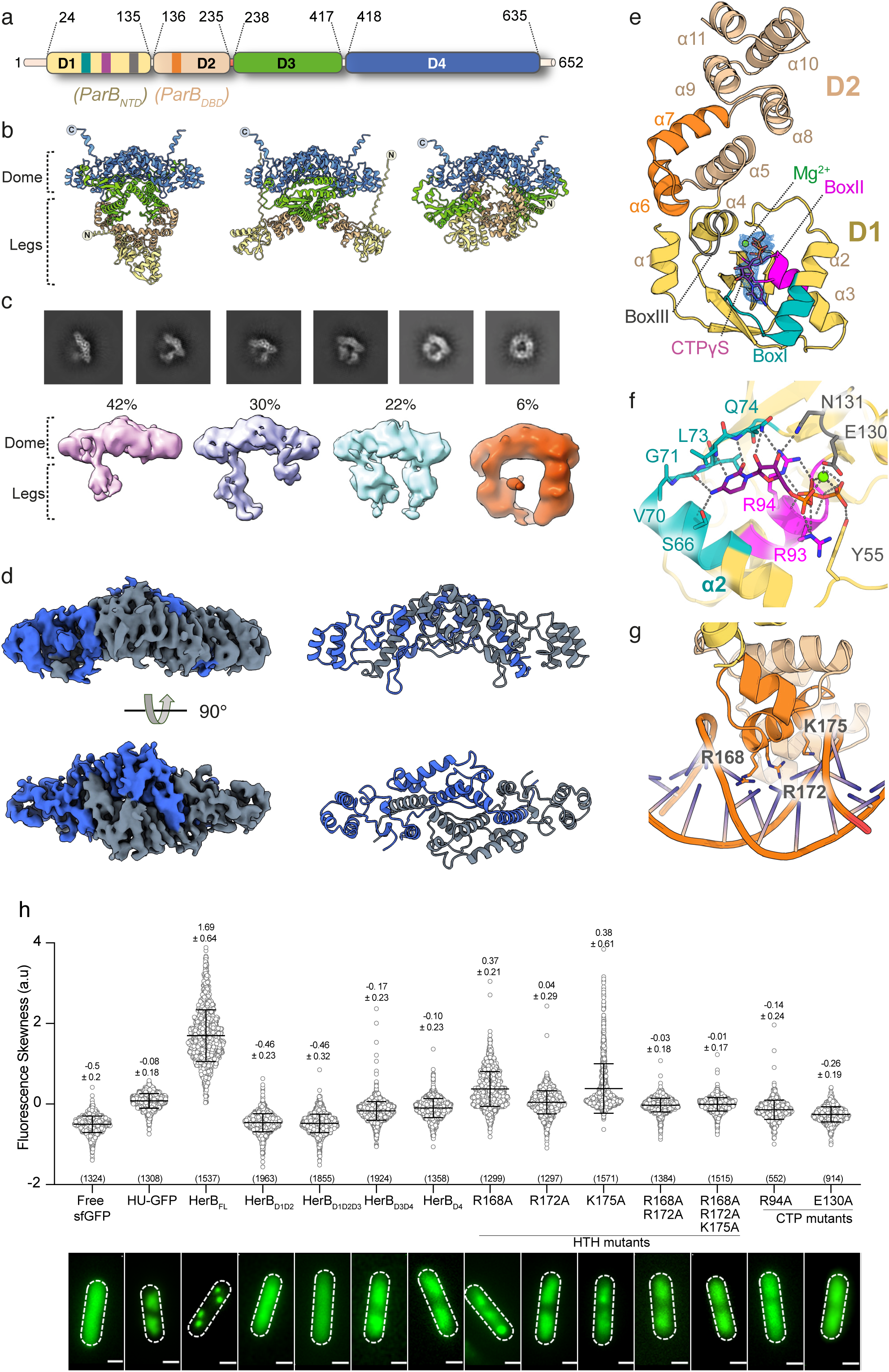
Structural definition of HerB. (**a**) Schematic representation of HerB domain boundaries identified by AlphaFold3 with the position of the CTP-binding motifs Box I (teal), Box II (magenta) and Box III (grey) and helix-turn-helix (orange) indicated. The corresponding ParB domains are indicated below. (**b**) Representation of three AlphaFold3 models of HerB dimer with domains coloured as in a). A “dome” is made of D4 dimers while legs consisting of D1, D2 and D3 are predicted with different orientations. (**c**) Representatives averaged-2D classes of HerB from CryoEM imaging. Below are represented the *ab initio* models of HerB with the corresponding percentage of particles used in each model. (**d**) Electron density map (contoured at level 0.152) of HerB_D4_ dimer (left) and the corresponding HerB_D4_ dimer structure (right), with chain A in blue and chain B in grey. (**e**) Crystal structure of HerB_D1D2_-CTPγS complex coloured according to a). 2Fo-Fc map is shown around the CTPγS and Mg^2+^ ion contoured at 1.5σ. (**f**) View of the CTPγS binding site with residues involved displayed as ball-and-stick. Polar contacts are represented by grey dashed lines. (**g**) Model of HerB_D1D2_ in complex with dsDNA, with side chains of DNA interacting residues displayed as ball-and-stick. (**h**) Jitter plot showing the skewness of intracellular green fluorescence in live *E. coli* cells expressing the indicated constructs, with representative microscopy images shown below (Scale bar, 1 μm). Grey circles represent individual-cell measurements. The number of cells analysed (n) from at least three independent biological replicates is indicated, together with the corresponding mean ± s.d.

To determine the oligomeric state of HerB, the purified protein was analysed by size-exclusion chromatography coupled to multi-angle light scattering. This analysis revealed a single species of ∼146 kDa, consistent with a homodimer (theoretical monomer mass 71.5 kDa; Extended Data Fig. 2a,b). To define HerB structural organization, we analysed the protein using AlphaFold3^20^, which predicted a modular architecture of four domains flanked by a flexible N-terminal extension (residues 1-23) and a C-terminal tail predicted to form a short α-helix (635-652). The N-terminal region (residues 1-235) comprises domain 1 (D1; residues 1-136), homologous to the ParB NTD, and domain 2 (D2; residues 137-235), corresponding to the ParB DBD containing a HTH motif^21,22^. The C-terminal half contains two additional domains, D3 (residues 238-417) and D4 (residues 418-652), with no detectable homology to previously characterized proteins (Fig. 2a). Notably, AlphaFold3 modelling of HerB dimer revealed that the two D4 domains assemble to form a V-shaped, dome-like core, while the remaining domains (D1-D3) extend outward as flexible “legs” capable of adopting various conformations (Fig. 2b, Supplementary Video S4, Extended Data Fig. 2c).

Cryo-EM analysis of vitrified full-length HerB revealed particles adopting multiple conformational states consistent with this architecture (Fig. 2c). Despite this heterogeneity, the V-shaped D4 dimerization core was readily identifiable, connected to one or two flexible “legs” and occasionally forming closed ring-like conformations. One particle subset yielded a 3.6 Å reconstruction, which improved upon application of two-fold symmetry during refinement (Fig. 2c, Extended Data Fig. 3a,b). This map enabled fitting and refinement of the D4 dimer model, except for the C-terminal extension (residues 637-652) predicted to form a helix (Fig. 2d; Extended Data Fig. 3c,d; Supplementary Table 1).

The cryo-EM structure of D4 (HerB_D4_; residues 421-636) reveals a helical fold organized into two subdomains (Fig. 2d, Supplementary Video S5). The first subdomain mediates tight dimerization, burying ∼4,500 Å² of surface area and involving domain swapping of an extended helix, differs from canonical ParB C-terminal domains^23,24^. Dimerization also brings β-strands from each monomer (amino acids 475-479) together to form a short intermolecular β-sheet. The second subdomain forms a small globular module of four short α-helices (residues 577-625) extending from the dimerization core. Interestingly, its surface contains a conserved basic patch and a protruding loop (residues 545-558) (Extended Data Fig. 4a,b).

To gain insights into the structure of the leg domains, we determined the crystal structure of HerB ParB-like domains D1-D2 bound to CTPγS (HerB_D1D2_-CTPγS) at 2.6 Å resolution (Fig. 2e,f; Supplementary Video S6; Supplementary Table 2). Residues 1-23 were not resolved, consistent with the flexible N-terminal segment predicted by AlphaFold3. The structure closely resembles canonical ParB proteins and contains one CTPγS molecule and a magnesium ion bound at the conserved nucleotide-binding pocket. D1 forms the CTP-binding module, whereas D2 comprises seven α-helices and contains the HTH motif formed by α6 and α7 (Fig. 2e; Extended Data Fig. 5a). CTPγS is coordinated by the conserved C motif (Box I), together with phosphate-binding motifs Box II (G91-A97) and Box III (M128-L132), which contact the nucleotide and Mg²⁺ ion via E130 (Fig. 2f, Extended Data Fig. 5a,b). Notably, Box III resides in a loop rather than an α-helix, and in contrast to previously characterized ParB structures, HerB_D1D2_-CTPγS crystallized as a monomer adopting a distinct conformation (Extended Data Fig. 5b,c). To assess whether this conformation could support DNA binding, we modelled a HerB_D1D2_-dsDNA complex by superposition with the *Caulobacter crescentus* ParB-*parS* structure^25^ (Extended Data Fig. 5e). This model positions the HerB HTH motif within the DNA major groove, with residues R168, R172 and K175 oriented to contact the DNA backbone and bases with no steric clash (Fig. 2g).

Together, these structural analyses reveal that HerB forms a modular dimer in which the novel D4 domain generates a stable dimerization core, while flexible ParB-like “legs” bearing CTPase and DNA-binding modules are positioned to engage chromosomal DNA.

### HerB focus formation requires DNA binding, CTP binding and D4

To determine the functional contribution of the structural domains identified above, we engineered targeted HerB mutants and analysed their effect on focus formation *in vivo*, a hallmark of ParB protein assembly^26^. Quantitative snapshot microscopy of exponentially growing cells expressing fluorescently tagged HerB variants was used to measure fluorescence skewness, an unbiased metric that increases as fluorescence becomes concentrated into discrete intracellular structures such as foci. Skewness values remain low when fluorescence is diffuse and increase when signal concentrates into discrete foci. As a reference for diffuse fluorescence, cells producing free fluorescent proteins (sfGFP) displayed low skewness values (−0.5 ± 0.2), reflecting the symmetric distribution of fluorescence throughout the cytoplasm (Fig. 2h). By contrast, nucleoid-associated HU-GFP, which does not form discrete foci but remains confined to the nucleoid region, exhibited significantly higher skewness values (−0.08 ± 0.18), reflecting asymmetry in fluorescence distribution despite the absence of focal structures (Fig. 2h). Remarkably, wild-type HerB-sfGFP fluorescence was strongly concentrated into bright intracellular foci with minimal diffuse signal, resulting in markedly higher skewness values (1.69 ± 0.64) (Fig. 2h).

Truncation analysis revealed that proteins lacking the C-terminal domains (HerB_D1D2_-sfGFP and HerB HerB_D1D2D3_-sfGFP) displayed diffuse cytoplasmic fluorescence with skewness values similar to free fluorescent proteins. In contrast, constructs containing the C-terminal domains D4 (HerB_D4_-sfGFP and HerB_D3D4_-sfGFP) showed fluorescence confined to the nucleoid region, with significantly increased skewness values, comparable to HU fusions (Fig. 2h). These results suggest that D4 mediates nucleoid association, whereas full-length HerB is required for discrete focus formation. Our modelling of HerB_D1D2_ interaction with dsDNA positioned residues R168, R172 and K175 of the HTH motif to engage the DNA major groove (Fig. 2h). Single substitutions within the motif (R168A, R172A and K175A), as well as double and triple mutants, strongly reduced focus formation while retaining nucleoid enrichment, indicating impaired higher-order assembly rather than loss of DNA association. Besides the HTH motif, CTP binding is known to control focus formation and assembly of ParB-family proteins^27–29^. We therefore examined the role of nucleotide binding in HerB. Substitution of Box II residue R94 (Fig. 2g), equivalent to *Bacillus subtilis* ParB R80 required for CTP binding (Soh *et al.*, 2019), abolished focus formation while retaining nucleoid enrichment (skewness −0.14 ± 0.24). Mutation of E130 (E130A) involved in Mg^2+^ coordination (Fig. 2g) and required for CTP hydrolysis in *C. crescentus* (E135)^25^ similarly affected formation of discrete foci (Fig. 2h).

Together, these results demonstrate that all of HerB domains, as well as functional HTH and CTP binding motifs are all required for focus formation, consistent with assembly of CTP-dependent DNA-associated dimeric complexes on the chromosome.

### HerB reprograms host transcription through chromosomal association

To investigate the biological function of HerB, we compared the transcriptomes and proteomes of cells ectopically expressing HerB from pTrc99a-*herB-sfGFP* with those of cells carrying the empty vector. RNA-seq revealed extensive transcriptional remodelling, with 505 genes significantly repressed and 361 induced (Fig. 3a). Among these were 52 transcriptional regulators, including global regulators such as *arcA*, *narL, rcsA* and *hcaR*, indicating that part of the response likely arises from secondary regulatory cascades (Extended Data Fig. 6a). For example, expression of the global regulator ArcA was reduced in HerB-producing cells (Log_2_Fold-change = −1.31), accompanied by reciprocal changes in ArcA-regulated genes (Supplementary Table 3). To focus on transcriptional effects more directly associated with HerB, genes belonging to these secondary regulons were considered as indirectly regulated (Fig. 3a). Functional enrichment analysis revealed coordinated modulation of pathways involved in anaerobic respiration, glycolysis and TCA cycle, heme and iron metal binding, branched-chain amino acid biosynthesis, sulfur and nitrate metabolism, indicating extensive metabolic rewiring (Fig. 3b; Supplementary Table 3). To determine whether these transcriptional changes propagated to the proteome, we performed Tandem Mass Tag (TMT) quantitative proteomics. Although fewer proteins were detected as significantly altered than transcripts, the direction of regulation was largely conserved across pathways (Extended Data Fig. 6b). This concordance provides independent support for HerB-driven metabolic reprogramming. Remarkably, transcriptomic analysis also revealed coordinated regulation of genes controlling growth-phase transitions and translational activity (Extended Data Fig. 6a). The anti-sigma factor gene *rsd*, which encodes an inhibitor of σ⁷⁰-dependent transcription and contributes to the shift toward stationary-phase gene expression, was strongly upregulated in HerB-producing cells. In parallel, the ribosome hibernation factor *rmf* and the ribosome-associated inhibitor A *raiA*, which normally inactivate ribosomes and reduce translation during growth arrest, were markedly repressed. Together, these regulatory changes suggest a physiological configuration in which remodelling toward the stationary phase transcriptional program occurs while ribosome activity is maintained.

**Figure 3.**
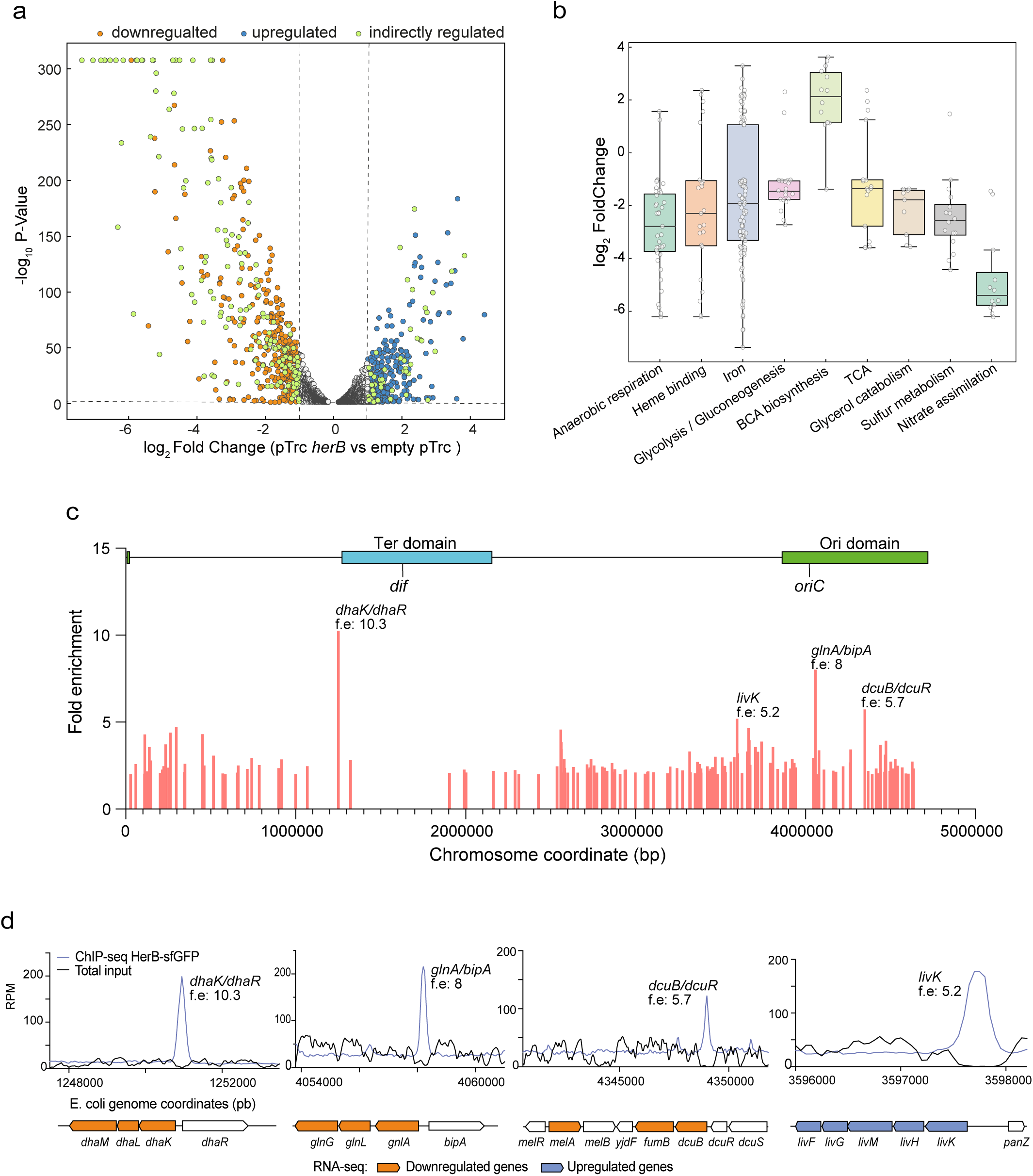
HerB forms intracellular foci, associates with the chromosome and reprograms host gene expression. (**a**) Volcano plot of RNA-seq results comparing cells carrying pTrc-*herB* with cells carrying the empty vector. Dashed lines indicate the log_2_ fold-change thresholds (−1 and +1) and the significance threshold (P = 0.05). Upregulated genes are shown in blue and downregulated genes in orange. Light-green dots indicate genes belonging to regulons controlled by transcriptional regulators whose expression is altered in the presence of HerB and are therefore considered indirectly regulated. (**b**) Plot showing the functional clusters identified using the DAVID bioinformatics resource (https://david.ncifcrf.gov/). The median and interquartile range (Q1–Q3) are indicated by the boxes, the mean by a horizontal black line, and the minima and maxima by the whiskers. Grey dots correspond to the log_2_ fold-change values of individual genes within each functional category. (**c**) Distribution and fold enrichment (f.e.) of HerB ChIP-seq peaks (total input-subtracted) during ectopic HerB production mapped onto the linearized *E. coli* chromosome. The boundaries of the Ori and Ter macrodomains are indicated above. (**d**) Genome-browser views showing the four most enriched HerB ChIP-seq peaks. Read density is represented as reads per million (rpm). The names of the genes associated with each peak and the corresponding fold enrichment (f.e.) are indicated. Schematics of the corresponding genomic regions are shown below. Genes significantly downregulated or upregulated in the RNA-seq dataset are coloured orange and blue, respectively.

Physiological assays confirmed that these transcriptional changes translated into coherent cellular phenotypes. Consistent with repression of anaerobic respiration genes, HerB-producing cells exhibited reduced resazurin reduction (Extended Data Fig. 7a), indicating altered redox metabolism. Induction of *yhjX* encoding a pyruvate transporter in RNA-seq data, suggested increased pyruvate excretion and was confirmed by elevated extracellular pyruvate levels (Extended Data Fig. 7b). Despite repression of *aceEF*, acetate production did not increase, indicating rerouting of carbon flux toward pyruvate accumulation (Extended Data Fig. 7c). Similarly, upregulation of fimbrial genes correlated with reduced motility (Extended Data Fig. 7d), demonstrating that transcriptional remodelling results in measurable physiological consequences.

To test whether these changes reflected direct chromosomal association, we performed ChIP-seq on cells ectopically expressing HerB-sfGFP (Fig. 3c). HerB was enriched at 156 chromosomal regions distributed all over the host chromosome, except for a noticeable depletion from the terminus region (Fig. 3c; Supplementary Table 3). The extent of HerB-enriched regions was highly variable, ranging from 346 to 2971 bp (median ∼532 bp). Among these 156 peaks, 71 peaks (45.5 %) occurred within genes or regulatory regions of operons displaying coordinated expression changes in the RNA-seq dataset and 4 occurred within prophages (Fig. 3d; Extended Data Fig. 8a). Together, these observations support a non-random relationship between HerB binding and transcriptional changes and suggest that binding at a single locus may influence entire operonic units. Furthermore, the broad and variable size of these regions indicates that HerB association may extend over contiguous chromosomal segments rather than being confined to discrete sequence-specific targets. Consistent with this observation, systematic searches for enriched sequence motifs within HerB-bound regions failed to identify a strong consensus sequence or inverted repeat reminiscent of canonical ParB binding sites, *parS*.

Because HerB-sfGFP is expressed specifically in newly formed transconjugants, ChIP-seq performed during conjugation selectively captures DNA fragments bound in this subpopulation. Analysis 1.5 hours after mating initiation identified 20 reproducible HerB-enriched chromosomal loci, all of which corresponded to peaks observed in HerB-sfGFP ectopic expression condition (Extended Data Fig. 8b), and 13 of which corresponded to genes differentially expressed in RNA-seq, 4 to prophages, and 3 to genes encoding regulatory proteins or RNAs (Supplementary Table 3). No enrichment was detected on the F plasmid itself. Additional peaks mapped to regulatory or stress-related regions, including the small RNAs *gcvB* and *sraG*, the stringent starvation protein *sspA*, and a locus within the KpLE2 phage-like element (Supplementary Table 3).

Together, these transcriptomic, proteomic and chromatin-association analyses indicate that HerB binds the chromosome and drives extensive transcriptional remodelling that reshapes host metabolism and physiology.

### HerB alleviates plasmid acquisition cost and facilitates plasmid dissemination

To determine the contribution of HerB to plasmid transfer and establishment, we first compared conjugation efficiencies of the wild-type F and F*ΔherB* plasmids. No significant differences in transconjugant frequencies were detected after 1, 2 or 3 h of conjugation, indicating that HerB deletion does not affect transfer frequency under our experimental conditions (Fig. 4a).

**Figure 4.**
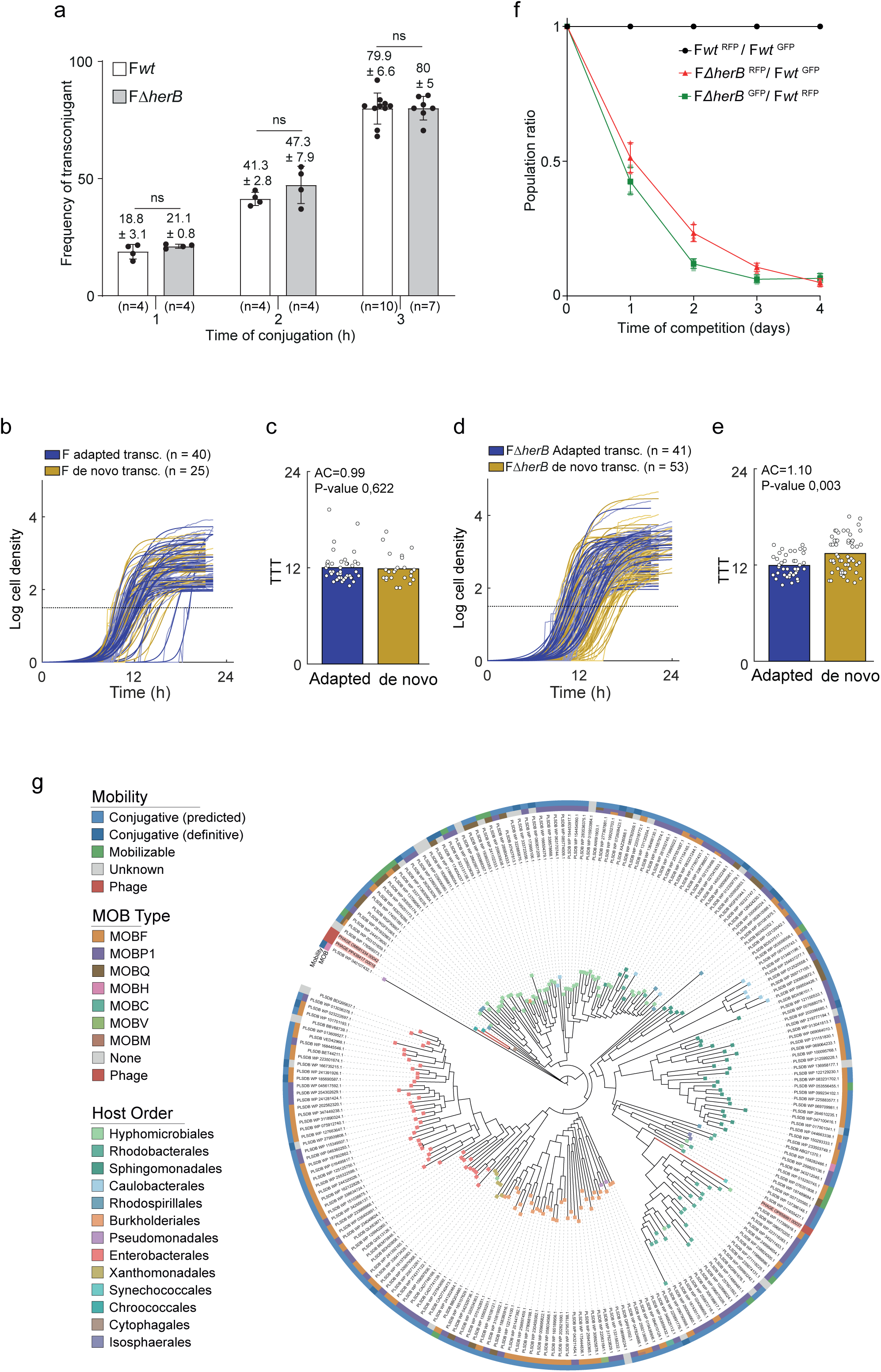
HerB alleviates acquisition cost, promotes plasmid dissemination and is widely conserved. **(a)** Frequency of transconjugants (100 x T/(R+T)) measured 1, 2 and 3 h after conjugation between recipient cells and donor cells carrying either the wild type F plasmid or F*△herB*. The mean with the corresponding ± s.d. obtained on (n) independent biological replicates is indicated, together with indivilual data points (black dots). *P*-value were calculated using One-way analysis of variance (ANOVA) followed by Dunnets multiple comparisons test (n.s., non-significant). **(b)** Growth trajectories of individual adapted and *de novo* transconjugant colonies following acquisition of the wild type F plasmid. Dashed line indicates the threshold used to calculate time-to-threshold (TTT=1.5). **(c)** Quantification of acquisition cost for wild type F plasmid acquisition, measured as changes in time-to-threshold between adapted (purple) and *de novo* (yellow) transconjugants. No significant acquisition cost was detected under these conditions (AC = 0.99, P = 0,622). **(d)** Growth trajectories of individual adapted and *de novo* transconjugants following acquisition of the F*△herB* plasmid. **(e)** Quantification of acquisition cost following F*△herB* acquisition. De novo transconjugants exhibited delayed time-to-threshold relative to adapted controls, corresponding to a significant acquisition cost (AC = 1.10, P = 0,003). Data in all panels represent individual colonies from three independent biological replicates. The total number of colonies analysed (n) is indicated and represents the sample size used for statistical analysis. **(f)** Plasmid dissemination competition assay showing the relative abundance of two donors carrying either the wild type F plasmid or F*△herB*, and producing either the green mChartreuse or the red mScarlet2 fluorescent protein. The conjugation mixture initially contained donor and recipient cells at a 1:1:5 ratio, and was diluted every 12 h into fresh medium containing recipient cells over a period of 4 days. The relative abundance of red- and green-fluorescent cells was quantified every 24 h by flow cytometry. **(g)** Maximum likelihood phylogenetic tree of 227 HerB homologues from plasmid (PLSDB) and phage (INPHARED) databases. Leaf colors indicate host order; inner ring shows MOB relaxase type; outer ring shows mobility classification. Three phage-associated homologues are highlighted in red.

The early and transient production of HerB in transconjugant cells prompted us to investigate whether it instead contributes to post-transfer events associated with plasmid establishment. Acquisition of a plasmid often imposes a transient physiological burden on recipient cells, referred to as the acquisition cost, which affects the early growth dynamics of newly formed transconjugants^30,31^. These acquisition costs can arise through changes in growth rate, lag time, or both, and are therefore reflected in altered time-to-threshold dynamics of newly formed transconjugants. To quantify this effect, we monitored the growth of individual colonies immediately after plasmid acquisition. Acquisition of the wild type F plasmid was not associated with any significant acquisition cost under our experimental conditions (Fig. 4b,c). In contrast, acquisition of the F*ΔherB* plasmid increased the acquisition cost by ∼10% (P-value 0,003), revealing that HerB contributes to optimal early growth of newly formed transconjugants (Fig. 4d,e).

We next asked whether this acquisition cost translated into a measurable disadvantage during plasmid dissemination. To address this, we developed a conjugation-competition assay to compare the spread of two plasmids competing for transfer into a recipient-cell population over several days. In this assay, two donor strains carrying plasmids marked with either red fluorescent mScarlet2 (F^RFP^) or green fluorescent mChartreuse (F^GFP^) were mixed with recipient cells at an initial 1:1:5 ratio (Fig. 4f; Extended Data Fig. 9a,b). Every 12 h, conjugation mixtures were diluted into fresh medium supplemented with naïve recipient cells, thereby generating repeated cycles of plasmid transfer over four days. The relative abundance of each plasmid lineage was quantified every 24 h by flow cytometry (Fig. 4f; Extended Data Fig. 9c). Compared with control competitions between isogenic wild-type plasmids, which served as a reference, direct competition between F*wt* and F*ΔherB* plasmids revealed a progressive decline of the F*ΔherB* plasmid in the population, irrespective of the fluorescent marker used. Together, these results demonstrate that although HerB does not measurably affect short-term conjugation efficiency (Fig. 4a), its ability to reduce acquisition costs in newly formed transconjugants (Fig. 4b-e) confers a substantial advantage during repeated cycles of plasmid transfer and establishment (Fig. 4f).

### HerB homologues are widespread among conjugative plasmids

To assess whether HerB is restricted to the F plasmid or more broadly distributed among mobile genetic elements, we performed an iterative profile HMM search against a dereplicated set of proteins from the PLSDB plasmid database^32^ and the INPHARED bacteriophage database^33^. Using HerB as a query, the iterative similarity searches identified 429 HerB homologues covering at least 80% of each of the four HerB functional domains, from which 227 representatives were selected after clustering at 70% sequence identity. The plasmid HerB homologues were encoded by a diverse set of conjugative and mobilizable plasmids encoding relaxases from multiple Mob families (MobF, MobP1, MobQ, MobH, MobC, MobV, and MobM), with hosts spanning Alpha-, Beta-, and Gammaproteobacteria, as well as a single representative from Bacteroidota and two from Planctomycetota (Fig. 4g). This distribution indicates that HerB-like proteins occur across genetically distinct plasmid backbones and conjugation systems. In addition to the plasmid-encoded homologues, three HerB-like proteins were detected in cyanophage genomes infecting *Synechococcus* and *Microcystis*, falling into two distinct clades in the protein tree. The functional significance of this association remains unclear.

Together, these observations indicate that HerB defines a widely distributed family of mobile-element proteins, suggesting that the mechanism described here extend well beyond F-like plasmids.

## Discussion

The post-entry phase of conjugation represents a critical bottleneck for successful plasmid establishment in a naïve host. Our results identify HerB as a transient, zygotically induced leading-region factor that operates during this window. Produced immediately after plasmid entry and persisting for only a few generations, HerB triggers extensive and structured remodelling of host gene expression, affecting central carbon metabolism, redox balance, biosynthetic pathways and motility. Importantly, this reprogramming is not merely a molecular signature: by alleviating acquisition cost in newly formed transconjugants, HerB confers a substantial advantage during repeated cycles of plasmid transfer and establishment. Thus, HerB links early host physiological remodelling to plasmid dissemination success.

A prominent component of this response is metabolic reconfiguration. Increased oxidative activity, repression of anaerobic fermentation pathways and elevated extracellular pyruvate indicate altered carbon partitioning and redox balance. These features are consistent with overflow metabolism, in which excess carbon flux is redirected through glycolysis and secreted as metabolites such as pyruvate^34,35^. In *E. coli*, overflow metabolism arises from proteome allocation constraints: although respiratory pathways are energetically efficient, they require greater proteomic investment than fermentative routes. As biosynthetic demand increases, cells shift toward metabolic strategies that reduce proteome investment while sustaining high rates of macromolecular synthesis^34,36^. Plasmid establishment likely imposes precisely such a demand. Following conjugative transfer, the incoming plasmid must be converted into double-stranded DNA while numerous plasmid-encoded proteins, particularly leading-region factors, are rapidly produced, placing a substantial burden on transcriptional and translational capacity. Within this framework, HerB-mediated transcriptional reprogramming may transiently shift host metabolism toward an overflow-like state, increasing glycolytic throughput and biosynthetic capacity while preventing metabolic bottlenecks and redox imbalance. The elevated extracellular pyruvate observed in HerB-producing cells is consistent with such regulated carbon overflow. Transcriptomic signatures further support this interpretation as *rsd* is upregulated, whereas the ribosome inactivation factors *rmf* and *raiA* are strongly repressed, a regulatory configuration expected to promote transcriptional remodeling while maintaining active ribosome pools and high translation capacity^37,38^. By promoting a metabolic state optimized for biosynthetic throughput rather than energetic efficiency, HerB may help recipient cells accommodate the intense demand for plasmid-encoded protein production immediately after transfer.

The structural architecture of HerB proposes a potential mechanism linking chromosome association to these global transcriptional effects. HerB combines ParB-like CTP-binding and helix-turn-helix DNA-binding modules with a previously uncharacterized C-terminal domain that mediates dimerization and nonspecific DNA association. ChIP-seq revealed HerB enrichment at multiple chromosomal loci distributed across the genome, consistent with direct DNA association *in vivo*. However, the breadth of the transcriptional response and the dispersed nature of HerB-enriched regions suggest that HerB does not act as a classical sequence-specific transcription factor. Instead, HerB chromosomal association probably contributes to transcriptional remodelling through a combination of direct local effects and indirect regulatory or metabolic cascades. The finite number of ChIP peaks, their variable width and the absence of a strong consensus motif or canonical *parS*-like inverted repeat suggest that HerB recruitment is not dictated by a simple DNA sequence code. Rather, HerB may preferentially associate with chromosomal regions defined by accessibility, local DNA topology, transcriptional activity or interactions with host factors.

This interpretation is consistent with the requirement for both CTP binding and the HTH motif for focus formation, together with the nucleoid-associated foci observed in living cells. These features support a CTP-dependent DNA-loading mechanism analogous to the clamp-like behaviour of ParB proteins. In this model, HerB dimers could load onto DNA directly or through an unidentified loading partner, and dynamically associate with extended chromosomal regions through sliding or spreading interactions. Such behaviour would explain the broad ChIP profiles, the absence of an identifiable binding motif and the ability of HerB to influence coordinated transcriptional programs across multiple metabolic pathways. The recovery of HerB foci after rifampicin washout further indicates that focus formation depends on higher-order nucleoid organization, rather than stable association with fixed intracellular sites alone. Although the molecular mechanisms by which HerB associates with DNA and modulates transcription remain to be defined, these features place HerB within the expanding family of bacterial CTP-dependent molecular switches harbouring a ParB-like CTPase fold^39^, while highlighting a function distinct from canonical segregation proteins.

Our findings broaden the conceptual scope of leading-region biology. Leading genes have primarily been described as anti-defence modules that protect incoming plasmids from restriction–modification systems, CRISPR-Cas immunity or SOS responses^4,5^. By contrast, HerB reveals that leading-region proteins can also promote plasmid success by actively reprogramming host physiology. In this model, zygotically produced HerB associates with the chromosome immediately after plasmid entry and initiates a coordinated transcriptional and metabolic program that accommodates the biosynthetic burden of plasmid establishment. This reprogramming alleviates acquisition cost, accelerates maturation of newly formed transconjugants and provides a competitive advantage during repeated transfer cycles.

HerB homologues are found in a broad range of conjugative and mobilizable plasmids spanning multiple Mob families and diverse bacterial genera, indicating that this strategy is not restricted to F-like plasmids. The broad phylogenetic distribution of HerB-like proteins suggests that transient host reprogramming may represent a recurrent solution evolved by mobile genetic elements to overcome the establishment bottleneck following horizontal transfer. More broadly, our work identifies host physiological subversion as a plasmid-encoded strategy that facilitates establishment and dissemination.

## Acknowledgements

The authors thank the National BioResource project, the Coli Genetic Stock Center, R. Reyes-Lamothe, J.Y. Bouet and J. Rech for providing strains, and C. Tessa for medium preparation. We thank the European Synchrotron Radiation Facility for providing access the CM01 cryo-EM and ID30B X-ray crystallography beamlines. **Funding**: This research was funded by the Foundation for Medical Research (grant number FRM-EQU202103012587 to C.L); the French National Research Agency (grant number ANR-18-CE35-0008, ANR-19-CE11-0012 and ANR-23-CE12-0037), the University of Lyon for funding to C.V., and the National Institutes of Health award (#1R35GM150871-01 to AJL). This work used the EM facilities at the Grenoble Instruct-ERIC Center (ISBG; UMS 3518 CNRS CEA-UGA-EMBL) with support from the French Infrastructure for Integrated Structural Biology (FRISBI; ANR-10-INSB-05-02) and GRAL, a project of the University Grenoble Alpes graduate school (Ecoles Universitaires de Recherche) CBH-EUR-GS (ANR-17-EURE-0003) within the Grenoble Partnership for Structural Biology. The IBS Electron Microscope facility is supported by the Auvergne Rhône-Alpes Region, the Fonds Feder, the Fondation pour la Recherche Médicale and GIS-IBiSA. **Author contributions:** C.V., S.F. and A-BD constructed strains and plasmids; C.V., S.F. and S.B. acquired and analysed microscopy images; C.V., S.F. and J.G. achieved protein purification and MALS; D.T. and L.T. prepared grids, collected and processed cryoEM data and analysed the cryoEM structure; C.V., G.P. and S.B. prepared and analysed Chip-seq samples; C.V., F.D. and A.P. did the mass-spec experiments. D.B. and Y.B. performed the phylogenetic analysis. AJL and SMA performed the fitness experiment. L.T. and C.L. wrote the paper and prepared the figures, with input from all authors. L.T., Y.Y., AJL, P.V. and C.L. hosted the research and provided funding.

## Methods

### Bacterial strains, plasmids and growth

Bacterial strains are listed in Supplementary Table 4, plasmids in Supplementary Table 5 and primers in Supplementary Table 6. Fusion of genes with fluorescent tags and gene deletion on the F plasmid used λRed recombination^40,41^. Modified F plasmids were transferred to the background strain K12 MG1655 by conjugation. When genetic modification on the F plasmid were required, the *kan* and *cat* gene were removed using site-specific recombination induced by expression of the Flp recombinase from plasmid pCP20^40^. Plasmid cloning was done by Gibson Assembly and verified by Sanger sequencing (Eurofins Genomics biotech). Cells were grown at 37°C in M9 medium supplemented with glucose (0.2%) (M9-Glucose) and casamino acid (0.4%) (M9-CASA) before imaging; Rich Defined Medium (RDM, Teknova) was also used for some microscopic experiments. When needed, glucose was replaced as carbon source by glycerol (0.2%) or maltose (0.2%). Cells were grown at 37°C in Luria-Bertani (LB) broth for other assays (Sigma-Aldrich). When appropriate, supplements were used in the following concentration: Ampicillin (Ap) 100 μg/mL, Chloremphenicol (Cm) 20 μg/mL, Kanamycin (Kn) 50 μg/mL, Streptomycin (St) 20 μg/mL and Tetracyclin (Tc) 10μg/mL.

### Conjugation assays

Overnight cultures in LB of recipient and donor cells were diluted to an A_600_ of 0.05 and grown until an A_600_ comprised between 0.7 and 0.9 was reached. 25 μL of donor and 75 μL or recipient cultures were mixed into an Eppendorf tube and incubated for 90 minutes at 37°C. 1 mL of LB was added gently and the tubes were incubated again for 90 minutes at 37°C. Conjugation mix were vortexed, serial diluted, and plated on LB agar X-gal 40 μg/mL IPTG 20 μM supplemented with the appropriate antibiotic to select for recipient or donor population. Recipient (R) colonies were then streaked on plates of LB agar containing Tetracyclin 10 μg/mL to select for transconjugants (T) and the frequency of transconjugant calculated from the (T/R+T) ratio presented in Figure 6A.

### Chromosome degradation

Strains carrying two *I-SceI^CS^*cut sites^18,19^ were transformed with the pTrc *herB-sfgfp* plasmid and the pSCL plasmid carrying the *I-SceI* gene under the control of the *P_ara_* promoter and plated on LB agarose plates containing 0.2% glucose, ampicillin Ap (100 μg/mL) and Cm (20 μg/mL) at 37°C. Transformant clones were propagated on LB agarose plates containing 0.2% glucose. Single colonies were inoculated in M9 CAA minimal medium supplemented with 0.2% glucose and Ap and incubated overnight at 37°C with agitation (140 rpm). The next day, overnight cultures were diluted (1/200) and grown to early exponential phase (OD_600nm_ ∼0.4). 0.2% arabinose was added to induce the production of I-SceI endonuclease and initiate chromosome degradation in the *recA-* strains, and cultures were incubated 120 min at 37°C for complete DNA degradation before microscopy.

### Live-cell microscopy

Overnight cultures in M9-CASA were diluted to an A_600_ of 0.05 and grown until A_600_ comprised between 0.7 and 0.9 was reached for conjugation analysis. Conjugation samples were obtained by mixing 25 μL of donor and 75 μL of recipient into an Eppendorf tube. For time-lapse experiments, 50 μL of the pure culture of conjugation mix was loaded into a B04A microfluidic chamber (ONIX, CellASIC®)^42^. Nutrient supply was maintained at 1 psi and the temperature maintained at 37°C throughout the imaging process. Cells were images every 1 or 5 minutes for 90 to 120 minutes. For snapshot imaging, overnight cultures in M9-CASA, M9-Glucose or RDM were diluted to an A_600_ and grown until A_600_ of 0.2 was reached for analysis of exponentially growing cells and 10 μL samples of clonal culture or conjugation mix were spotted onto an M9-CASA 1% agarose pad on a slide^43^ and imaged directly.

#### Image acquisition

Conventional wide-field fluorescence microscopy imaging was carried out on an Eclipse Ti2-E microscope (Nikon), equipped with x100/1.45 oil Plan Apo Lambda phase objective, ORCA-Fusion digital CMOS camera (Hamamatsu), and using NIS software for image acquisition. Acquisition was performed using 50% power of a Fluo LED Spectra X light source at 488 nm and 560 nm excitation wavelength. Exposure settings were 100 ms for Ypet, sfGFP and mCherry and 40 ms for phase contrast.

#### Image analysis

Quantitative image analysis was done using Fiji software with MicrobeJ plugin ^44^. For snapshot analysis, cells’ outline detection was performed automatically using MicrobeJ and verified using the Manual-editing interface. For time-lapse experiments, detection of cells was done semi-automatedly using the Manual-editing interface, which allows to select the cells to be monitored and automatically detect the cell outlines. Within conjugation populations, donor (no mCh-ParB signal), recipient (diffuse mCh-ParB signal), or transconjugant (mCh-ParB foci) categories were assigned using the “Type” option of MicrobeJ. Recipient cells were detected on the basis of the presence of red fluorescence above the cells’ autofluorescence background level detected in the donors. Among these recipient cells, transconjugants were identified by running MicrobeJ automated detection of the ParB fluorescence foci (Maxima detection). This approach was used independently of the presence or absence of the Ssb-Ypet or HerB-sfGFP fusions within donor and recipient cells. Within the different cell types, skewness, Signal/Noise Ratio (SNR) or cell length (μm) parameters were automatically extracted and plotted using MicrobeJ. SNR corresponds to the ratio (mean intracellular signal / mean noise signal), where the mean intracellular signal is the fluorescence signal per cell area and the noise is the signal measured outside the cells (due to the fluorescence emitted by the surrounding medium). By contrast with the total amount of fluorescence per cell, which is depending on the cell size/age and accounts for the background, SNR quantitative estimate is more appropriate for unbiased quantification of intracellular fluorescence over time. HerB-sfGFP and HerB-mCh foci were detected using MicrobeJ Maxima detection function, and foci localisation and fluorescence intensity were extracted and plotted automatically.

### Gene cloning, protein expression and purification

HerB and HerB_D1D2_ (residues 1-236) production plasmids were generated by inserting corresponding PCR products into pET-151D vector using a commercial kit (Invitrogen). The resulting HerB proteins were with a N-terminal 6 his tag and TEV enzymatic cleavage site. Proteins were produced and purified using the same protocol. Expression vectors were transformed into *E. coli* BL21 Star cells (Invitrogen). Cells were grown in 1 L flasks at 37°C in LB medium supplemented with Ampicillin 100 μg/mL until an A_600_ of 0.6 was reached. Protein expression was induced with 1 mM of IPTG during 16 hours at 20°C. Bacterial cells were centrifuged at 6000 g and dry pellets stored at −80°C. Harvested wells were resuspended in buffer A (25 mM NaPi pH 8, 300 mM NaCl, 5% Glycerol (v/v)) implemented with lysozyme 1 mg/mL (Sigma-Aldrich), 30 U/mL DNase I (Sigma-Aldrich) and anti-protease EDTA-free cocktail (Roche). The cells were lysed by sonication and debris removed by centrifuged at 20,000 g for 40 minutes at 4°C. Recombinant protein was purified by chromatography using a Nickel-loaded Hitrap Chelating HP column (Cytiva). Unbound material was extensively washed using 10 column volumes of buffer A. An additional washing step with 2 column volumes of buffer A supplemented with 1 M NaCl was done before elution of the protein with a linear gradient of buffer A supplemented with 500 mM gradient of imidazole over 8 column volumes. Peak fractions were pooled and the His tag was cleaved with TEV protease (500 μg/20 mg of eluted protein) in presence of 1 mM DTT and 0.5 mM EDTA in overnight dialysis in buffer A. HerB and HerB_D1D2_ were further purified by size exclusion chromatography (Superdex S200 increase 16/600, Cytiva) equilibrated in buffer A. Purity of the samples was assessed by SDS-PAGE. Freshly purified HerB and HerB_D1D2_ were concentrated to 10 mg/mL on 30 kDa and 3 kDa Amicon Ultra concentrators (Millipore), respectively and then stored at −80°C.

### Size exclusion chromatography-Multi-angle light scattering (MALS)

Size exclusion chromatography experiments coupled to multi-angle laser light scattering (MALS) and refractometry (RI) were performed on a superdex S200 10/150 GL (GE Healthcare) equilibrated in buffer 25 mM NaPi pH 8, 300mM NaCl, 5% Glycerol. 50 μL of proteins were injected at a concentration of 10 mg/mL. On-line MALS detection was performed with a mini DAWN-TREOS detector (Wyatt Technology Corp., Santa Barbara, CA) using a laser emitting at 690 nm and by refractive index measurement using an Optilab T-rex system (Wyatt Technology Corp., Santa Barbara, CA). Weight averaged molar mass (Mw) were calculated using the ASTRA software (Wyatt Technology Corp., Santa Barbara, CA).

### CryoEM data collection, processing, model building and refinement

The vitreous grids of HerB were prepared on Quantifoil R 1.2/1.3 holey carbon grids using a Vitrobot. The glow-discharged grids were mounted in the sample chamber at 8°C and 95% relative humidity, blotted, and plunge-frozen in liquid ethane at temperature −172°C. The frozen grids were tested on a Glacios transmission electron microscope (TEM) at IBS, and the grid preparation conditions were optimized in cycles. The final optimized grids were prepared using 5 μL of HerB sample at a concentration of 1.2 mg/mL spotted on Quantifoil R 1.2/1.3 holey carbon grids, incubated on the grid for 30 s, and back blotted for 12-14 s using two pieces of Whatman® Grade 1 filter paper. High-resolution dataset was collected at ESRF-Grenoble CM01 facility using a 300 kV Titan KRIOS TEM equipped with a Gatan K3 Summit direct electron detector and a Gatan energy filter. The data collection was automated using EPU version 2.5 (Thermo Fisher Scientific). Electron movies were collected in the counting mode at a nominal magnification of 105,000x. The total exposure time was 6 s with a total dose of 50 e^-^/Å^2^ and the movies were recorded as gain corrected MRC files. Images for HerB sample were collected using EPU software version 2.6.1 (Thermo Fisher Scientific) in the counting mode again at a nominal magnification of 105,000x yielding a pixel size of 0.84 Å. The exposure time was 5 s for each movie, accumulating to a total dosage of 50 e^-^/Å^2^. The beam-image shift was applied during data collection to increase data throughput. 8,396 movies were recorded as compressed MRC files. For both data collections, the energy filter was used with a slit width of 20 eV.

Data processing was carried out using Cryosparc ^45^. All frames in individual movies were aligned and contrast-transfer-function (CTF) estimations were performed using CTFFIND-4^46^ An initial automated picking procedure was applied to a set of 500 images and particles were classified into 50 classes. The 12 best classes were used as the template for picking the particles from the complete dataset. 7,380,495 particles were extracted with a 300-pixel box size and submitted to several rounds of 2D classification, resulting in 455,865 selected particles. These were used to ab initio building of 4 models that showed different maps with low resolution (Extended Data Fig.3a). Heterogenous refinement was used to refined and re-classified particles leading to one map with resolution of 4.7 Å map with 191,728 particles. This set of particles was further sorted to reach a number of 186,180 and used in non-uniform refinement ^47^ leading a 3.6 Å resolution map applying a C2 symmetry (Extended Data Fig.3a,b,c). This map corresponded remarkably to the dimer of D4 predicted by AF3. Fitting of the predicted domain was performed in ChimeraX^48^ and manually improved in Coot^49^. The final model was then subjected to real-space refinement in Phenix^50^ (Extended Data Fig.3c,d). Model validation was carried out with the Comprehensive validation tool from the Phenix package (Supplementary Table 1). Analysis of the surface accessible conserved residues on the HerB_D4_ dimer was performed with the Consurf server and default parameters^51^. The coordinates and electron density map have been deposited to the Protein Data Bank, under the accession number 29HU.

### Crystallization, structure determination and refinement

Crystals of HerB_D1-2_-CTPγS were obtained with the sitting drop vapor diffusion method using a Mosquito robot and MRC well crystallization plates (Molecular Dimensions). Drops consisting of 200 nL of protein (10 mg/mL), 1 mM MgCl_2_ and 1 mM CTPγS (Jena Biosciences) and 200 nL of reservoir solution were left at 19°C for two days. Crystals of HerB_D1-2_-CTPγS appeared in condition H7 of PACT Premier screen (Molecular Dimensions) with a reservoir solution consisting of 0.2 M Sodium acetate trihydrate, 0.1 M Bis-Tris propane pH 8.5 and 20% w/v PEG 3350. Crystals were flash frozen in reservoir solution supplemented with 1 mM CTPγS and glycerol 15% (v/v). Analysis of the likely composition of the asymmetric unit suggested that it contain one copy of HerB_D1-2_ subunit. Data were collected at the ID30B beamline of the ESRF and processed with XDS^52^ and AIMLESS^53^ from CCP4 program suite^54^. The crystal of HerB_D1-2_-CTPγS diffracted to a resolution of 2.6 Å and belonged to the space group P2_1_ (Supplementary Table 2). The structure was solved by molecular replacement using the AF2 prediction (Jumper et al., 2021) as a probe in PHASER with a separated search for D1 and D2 domains. The model of HerB_D1-2_-CTPγS was refined with final R_work_/R_free_ of 0.196/0.223 with excellent geometry (Supplementary Table 2) and the coordinates and structure factors were deposited in the Protein Data Bank with accession code 29HV.

### Chromatin immunoprecipitation sequencing ChIP-seq

#### Samples preparation

Cells were grown overnight in LB at 37°C and diluted at an A_600_ of 0.05 in 5 mL tubes, and incubated at 37°C with agitation until they reached an A_600_ comprised between 0.7 and 0.8. Conjugation mix of 10 mL of donor and 30 mL of recipient were done in 50 mL Falcon tube pre-heated at 37°C, and briefly vortexing before incubation at 37°C during 2 hours. 100 μL were taken to evaluate by serial dilution the conjugation efficiency (spread 50 μl of 10^-4^ and 10^-5^ dilution on LB implemented with Xgal 40 μg/ml, IPTG 20μM and LB implemented with Xgal 40 μg/ml, IPTG 20 μM, Tc10 μg/ml.

#### Formaldehyde treatment

400μL of a 1 mM Na phosphate (pH 8) and 1.08 ml of 37% formaldehyde were added to 40 mL of bacterial culture. Tubes were gently mixed and incubated 10 minutes at room temperature, and then at 30 minutes on ice. The cells were centrifugated at 5000 rpm for 15 minutes at 4°C (Bucket 3655 SORVAL ST 16R). The supernatant was discarded and the pellet washed twice with 20 ml of cold 1X PBS, pH 7.4. A third wash was done 1 ml of 1x PBS transferred to an Eppendorf tube before centrifugation at 8500 rpm (AcuSpin micro17R-eppendorf). The pellets were stored at −80°C for maximum 4 weeks.

### Transcriptomic experiment

#### Samples preparation

Cells were grown overnight in LB at 37°C and diluted at an A_600_ of 0.03 in 500 mL flasks, and incubated at 37°C with agitation until they reached an A_600_ comprised between 0.7 and 0.8. The equivalent number of cells as maximum 30 mg at A_600_ of 1 was harvested and washed in 100 μL of lysis buffer (10mM TrisHCl, 1mM EDTA pH 8, 1 mg/mL lysozyme). The RNA extraction was done according to the Nucleospin RNA plus kit directions (Macherey-Nagel). RNA concentrations were quantified using Qubit (Thermo Fisher Scientific) and Nanodrop (Thermo Fisher Scientific) devices and RNA integrity was verified by 1% agarose gel migration. Samples were sent to Montpellier GenomiX (MGX).

#### Data analysis

Data were given by MGX in Excel tables from which a more detailed analysis was performed. Significative data were selected by keeping only those with a Log_2_ Fold Change ≥ 1 or ≤ −1 and a -Log_10_ *P-Value* ≥ 1.3. Volcano plots were generated using InstantClue software^55^. Metabolic pathways were obtained by GOTerm analysis of their gene ID using the DAVID Bioinformatic Resources website (https://david.ncifcrf.gov/).

### Mass spectrometry experiments

#### Samples preparation

Same initial cells as the one for transcriptomic experiment were used. Cells were grown overnight in LB at 37°C and diluted at an A_600_ of 0.03 in 500 mL flasks, and incubated at 37°C with agitation until they reached an A_600_ comprised between 0.7 and 0.8. The equivalent number of cells as 10 mL at A_600_ 0.8 was harvested and washed twice with 1 mL of cold 50mM Tris pH 7.5. Pellets were kept at −80°C. After resuspension in 1 mL 50 mM Tris pH 7.5, cells disruption was carried out using TissueLyser II (QIAGEN) and 0.1 mm glass beads (Sigma-Aldrich). 1% Foscholine was added and the samples were incubated 90 minutes at 4°C on a rotating wheel. Cell debris was removed by ultracentrifugation for 1 hour at 4°C at 15 000 g. Supernatants were transferred to Eppendorf Protein Low Binding tubes and proteins were quantifies using BCA Protein Assay Kit (Pierce). Further analysis and data collection were done according to the same protocol used by Nolivos *et al*., 2019^16^. Volcano plot were generated using InstantClue software^55^, and association between genes was done using the String database (https://string-db.org/).

### Determination of extracellular pyruvate and acetate concentrations and AlamarBlue reduction

Cells were grown overnight in LB at 37°C with the appropriate antibiotic and diluted at an A_600_ of 0.05, and incubated at 37°C with agitation until they reached an A_600_ of 0.5 for pyruvate and acetate assays and 0.2 for AlamarBlue redox assay (Thermo Fisher Scientific). Samples were prepared according to the Pyruvate Assay or Acetate Assay kit directions (Sigma-Aldrich) to determine the pyruvate concentration in culture supernatant. These kits are based on pyruvate oxidase, acetate oxidase or alamarBlue reduction and data were measured automatically using the TECAN Spark multimode plate reader.

### Acquisition cost assay

Acquisition costs were quantified using a modified single-colony time-to-threshold (TTT) assay as previously described^30^. Briefly, donor, recipient, and adapted transconjugant strains were grown overnight from single colonies in 2 mL LB broth with appropriate antibiotics as needed. Cultures were pelleted, resuspended 1:1 in M9CA media supplemented with 0.4% w/v Glucose, and incubated for 5 min at room temperature. Donor and recipient cultures were then mixed 1:1 and incubated for 1 h at 25C to generate *de novo* transconjugants. Adapted transconjugants carrying the same plasmid were maintained under identical conditions in parallel. Following incubation, *de novo* and adapted populations were diluted and plated onto dual-selective agar plates at densities yielding isolated single colonies. Plates were placed on a flatbed scanner at 37C and imaged every 15 min for 24 h. Colony growth curves were extracted using automated image analysis (https://github.com/ajlopatkin/acquisition_cost_image_analysis), and TTT was defined as the time required for an individual colony to reach a fixed threshold density of 1.5 during exponential growth. Acquisition cost (AC) was quantified as the ratio between the average TTT of de novo transconjugants and the average TTT of adapted transconjugants. Statistical significance between adapted and *de novo* populations was determined using two-sided t-tests on TTT values obtained from three independent biological replicates.

### Conjugation-competition assay

Plasmid conjugation-competition assay allows comparing the relative dissemination of two plasmids producing either the green mChartreuse or the red mScarlet2 fluorescent protein. Donor and recipient cells were grown overnight in LB at 37°C. The conjugation mixture was initially prepared by mixing equal proportions of the two plasmid donors mixed with an excess of recipient cells, corresponding to D1:D2:R ratio of 1:1:5, in 5 ml of LB at 37°C. Every 12 hours over a period of 4 days, the conjugation mixture was diluted to a final OD_600_∼0.0008 into 5 ml of fresh medium containing recipient cells at OD_600_∼0.004. Every 24 hours, the relative abundance of red- and green-fluorescent cells was quantified by flow cytometry using an Attune NxT Acoustic Focusing Cytometer (Thermo Fisher Scientific). For each sample, 100,000 cells were analysed by acquiring the RFP (YL2) and GFP (BL1) signals. Gate analysis allows for the identification of each cell populations depending on their fluorescence signal, calibrated on the individual profiles of each donor and recipient clonal populations.

### HerB homologues identification and phylogenetic analysis

HerB homologues were identified by iterative profile HMM searches using jackhmmer (HMMER v3.4^56^ with the F plasmid HerB sequence (UniProt Q9S4W2) as the query. Searches were performed with a maximum of 10 iterations, a reporting E-value threshold of 10⁻⁸ and an inclusion threshold of 10⁻¹⁰ (-N 10 -E 1e-8 --domE 1e-8 --incE 1e-10 --incdomE 1e-10). The target database was constructed by combining proteins from PLSDB (v2025.11.24) ^32^ and INPHARED (v2025.08.31)^33^, each dereplicated independently using CD-HIT (v4.8.1)^57^ at 90% sequence identity with 80% coverage (-c 0.9 -s 0.8 -g 1). Hits were filtered to retain only sequences with ≥80% coverage of each of the four HerB functional domains (Domain I: residues 14-136; Domain II: residues 137-235; Domain III: residues 238-417; Domain IV: residues 418-632), yielding 429 high-confidence homologues (426 plasmid, 3 phage). Representative sequences were selected by clustering at 70% sequence identity and 80% length coverage using CD-HIT (-c 0.7 -s 0.8 -g 1), resulting in a final set of 227 sequences (224 plasmids, 3 phages).

Multiple sequence alignment was performed using MAFFT (v7.475)^58^ with the L-INS-i algorithm (--localpair --maxiterate 1000). Maximum likelihood phylogenetic reconstruction was performed using RAxML (v8.2.12)^59^ under the LG+Gamma+F model (PROTGAMMALGF) with 100 rapid bootstrap replicates (-f a -N 100) and midpoint rooting. The tree was visualized using iTOL (v6)^60^.

### Statistical analysis

*P*-value significance was analysed running specific statistical tests on the GraphPad Prism software. Single-cell data from quantitative microscopy analysis were extracted from the MicrobeJ interface and transferred to GraphPad. *P*-value significance of single-cell quantitative data was performed using unpaired non-parametric Mann-Whitney statistical test, which allows to compare differences between independent data groups without normal distribution assumption. *P*-value significance for the frequency of transconjugants obtained by plating assays were evaluated using One-way analysis of variance (ANOVA) with Dunnets multiple comparisons test, which allows to determine the statistical significance of differences observed between the means of three or more independent experimental groups against a control group mean (corresponding to the F *wt*). When required, *P*-value and significance are indicated on the figure panels and within the corresponding legend.

