## Supplementary information for "Plasmid-encoded host reprogramming promotes plasmid dissemination"

**This file includes:**

Extended Data Figures 1 to 9 with corresponding figure legends

Supplementary Tables 1 to 7 (Expect Supplementary Tables 3 submitted as an excel file)

Supplementary Video legends S1-S6

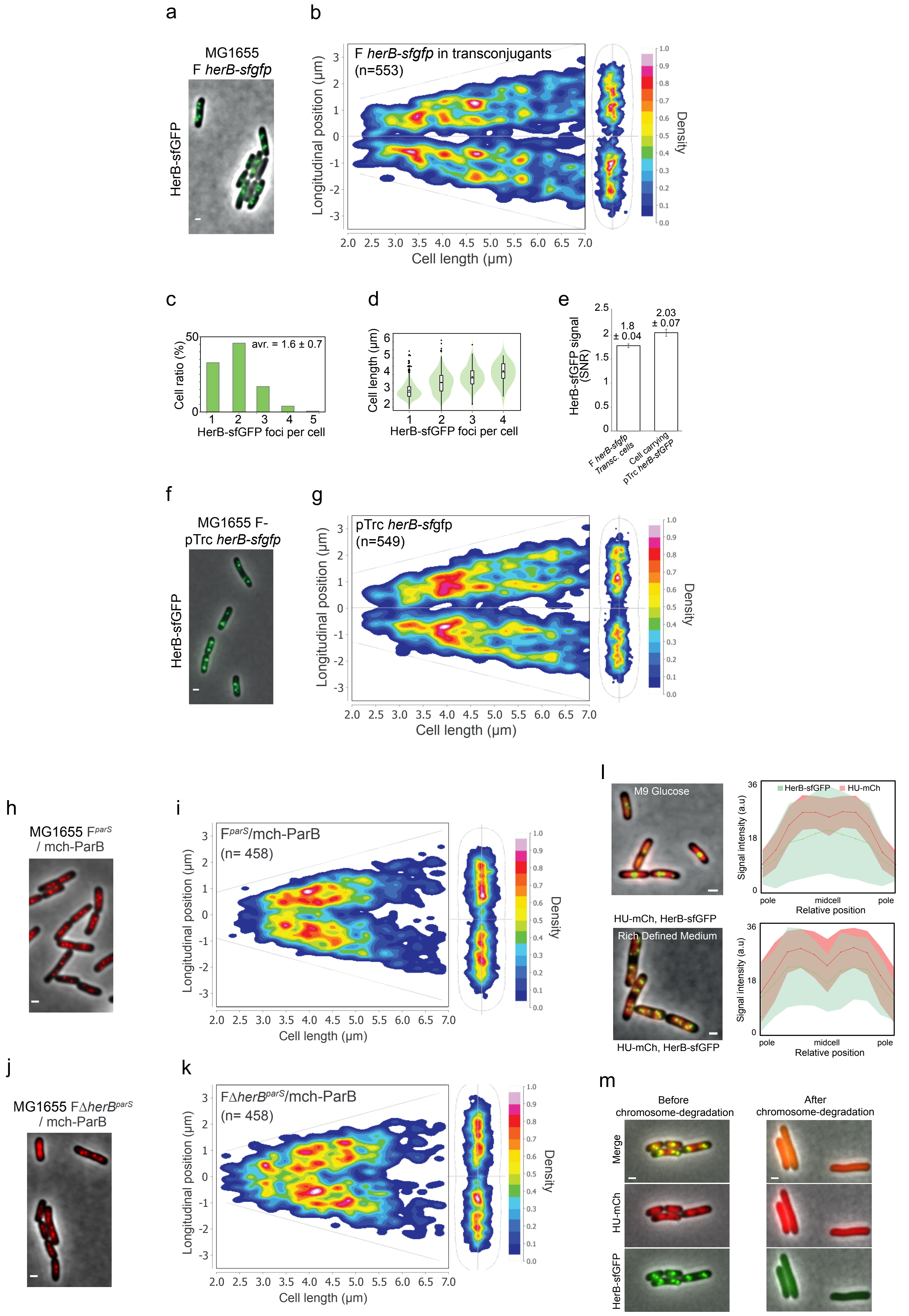

Extended Data Fig. 1

**Extended Data Figure 1.**

(**a**) Microscopy image of HerB-sfGFP produced from the F *herB-sfGFP* plasmid in transconjugant cells. Scale bar 1µm. (**b**) Localisation density maps of HerB-sfGFP in transconjugant cells having acquired the F *herB-sfGFP*. Localisation density in shown as a function of cell length (left) and within the outline of the normalised cell population (right). (**c**) Histogram of the proportion of transconjugant cells containing 1, 2, 3, 4 or 5 HerB-sfGFP foci. (**d**) Violin plot of cell length of transconjugants with 1, 2, 3 or 4 HerB-sfGFP foci. (**e**) Histogram of intracellular HerB-sfGFP fluorescence (signal-to-noise ratio) quantified in transconjugant cells or in cell carrying the ectopic pTrc99a *herb-sfGFP* plasmid. (**f**) Microscopy image of HerB-sfGFP ectopically produced from the pTrc99a *herb-sfGFP* plasmid. Scale bar 1µm. (**g**) Localisation density maps of HerB-sfGFP foci localisation in vegetative cells harbouring the ectopic pTrc99a *herB-sfGFP* plasmid. Localisation density in shown as a function of cell length (left) and within the outline of the normalised cell population (right). (**h**) Microscopy image showing the localisation or the F plasmid revealed by mCh-ParB/*parS* foci. (**i**) Localisation density maps of F plasmid positioning. (**j**) Microscopy image showing the localisation or the F*ΔherB* plasmid revealed by mCh-ParB/*parS* foci. (**k**) Localisation density maps of F*ΔherB* plasmid positioning. (**l**) Microscopy image of cells producing HU-mCh from the chromosome and HerB-sfGFP from the pTrc99a *herB-sfGFP* plasmid in cells grown in M9 Glucose or RDM rich defined medium. Scale bar 1µm. The quantitative analysis of HU-mCh and HerB-sfGFP fluorescence distributions along the normalised long axis of cells is presented in the left panels. Lines represent the mean, and the coloured area represent the standard deviation (s.d.). (**m**) Microscopy image of cells producing Hu-mCh from the chromosome and HerB-sfGFP from the pTrc99a *herB-sfGFP* plasmid. Degradation of the chromosome by induction of the I-SceI endonuclease in the absence of RecA triggers the loss of HU-mCh nucleoid structures as well as HerB-sfGFP foci.

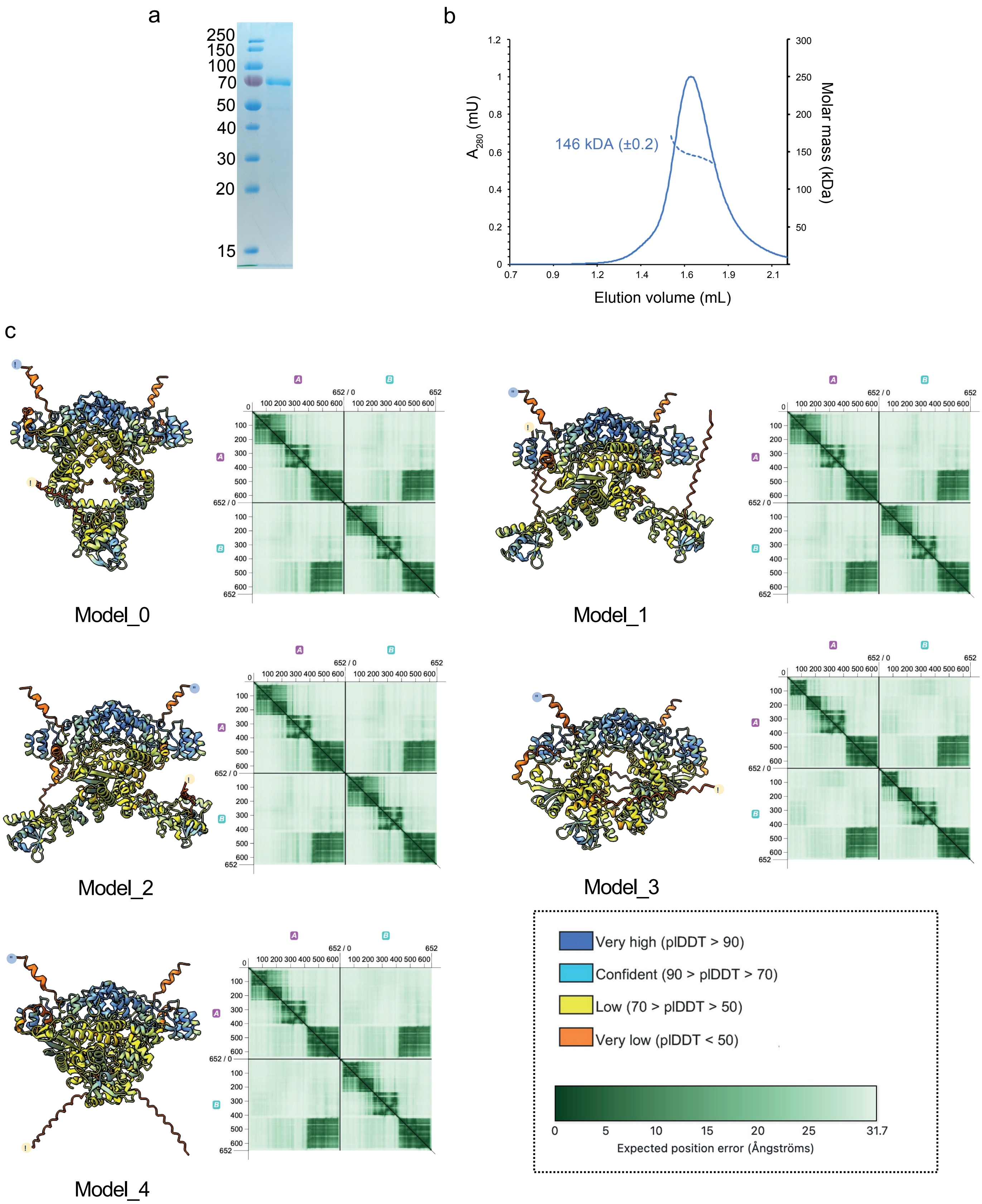

Extended Data Fig. 2

**Extended Data Figure 2. HerB dimer analysis**

(**a**) Coomassie-blue stained SDS-PAGE of purified HerB. (**b)** Size exclusion coupled to multi-angle light scattering measurement (MALS) of HerB. Chromatogram ($A_{280}$ absorption) of HerB on a Superdex 200 Increase 5/150 GL column. Dashed line indicates MALS calculation of the molar mass. (**c**) Five AlphaFold 3 models of HerB dimer (iPTM = 0.39, pTM = 0.41) with residues coloured according to their pLDDT and the corresponding predicted alignment error (PAE) matrices is shown.

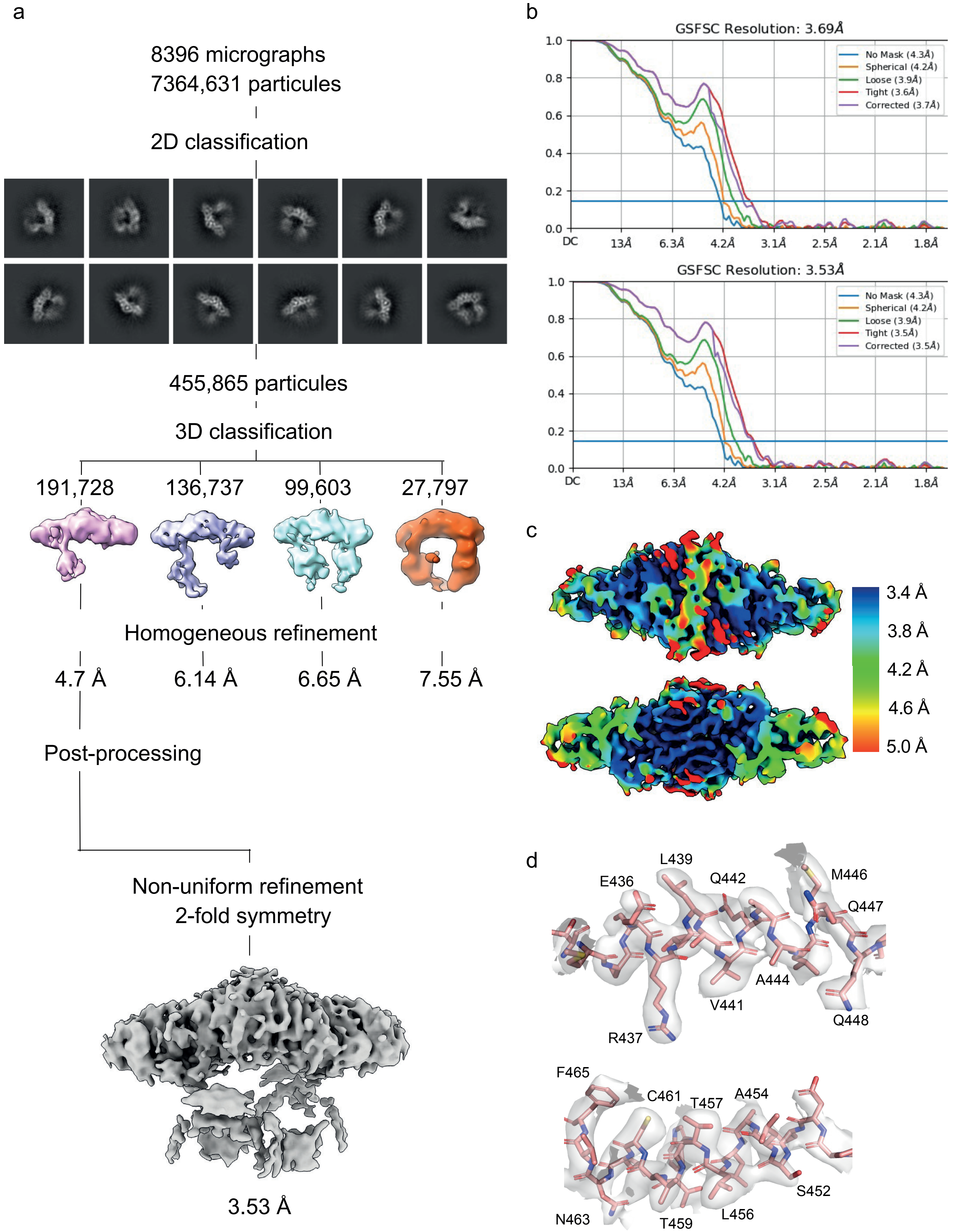

Extended Data Fig. 3

**Extended Data Figure 3. CryoEM structure determination of HerB$_{D4}$**

(**a**) Cryo-EM Data processing summary using Cryosparc. (**b**) Fourrier Shell Correlation curve for the final 3D reconstruction in Cryosparc. (**c**) Local resolution assessment of the final cryo-EM map. (**d**) Two images of HerB$_{D4}$ electron density (contoured at level 0.152) with model fitted displayed as balls-and-sticks.

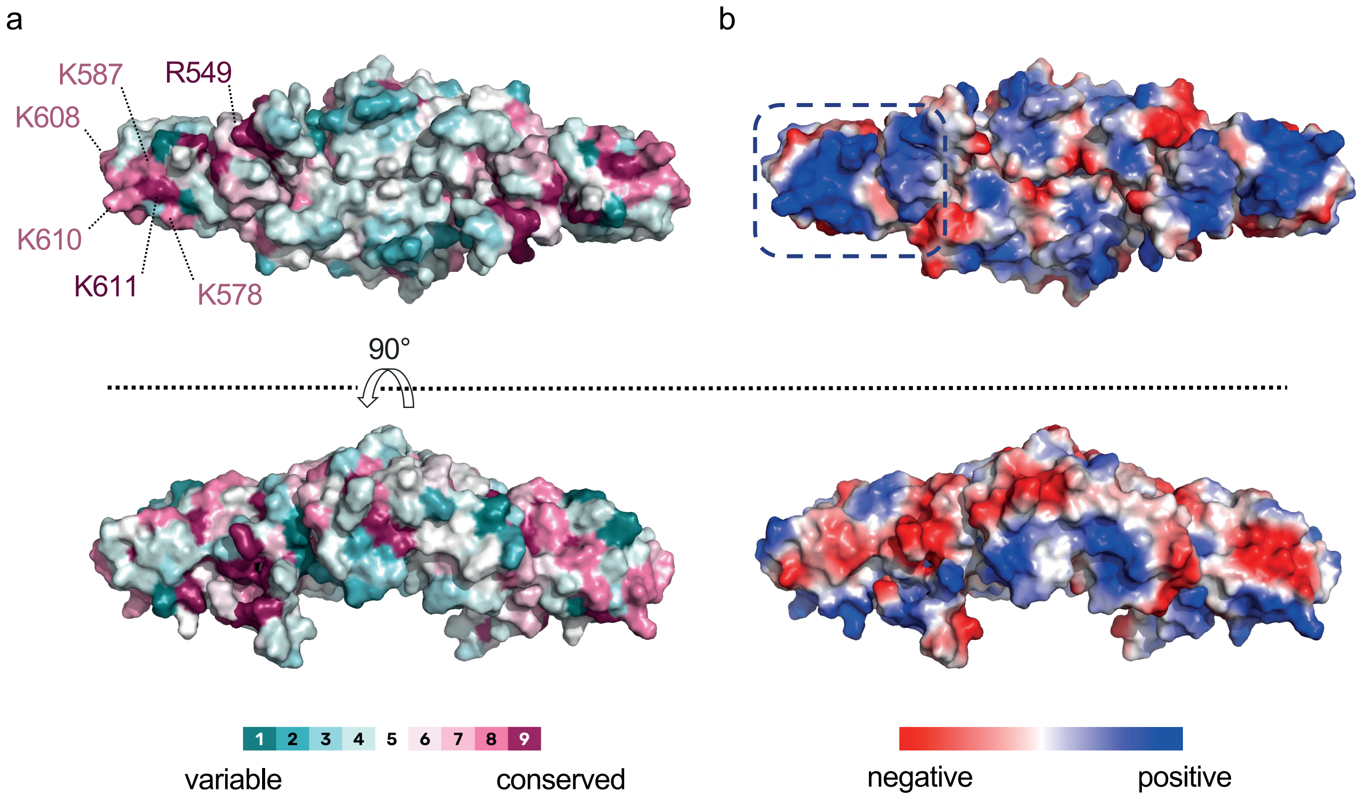

Extended Data Fig. 4

**Extended Data Figure 4. Structural analysis of $HerB_{D4}$ surface properties**

(**a**) Surface representation of $HerB_{D4}$ structure coloured according to sequence conservation. A patch of conserved residues is labelled. (**b**) Surface representation of $HerB_{D4}$ structure coloured according to electrostatic potential. The dashed line delineates the basic patch formed by conserved and positively charged residues labelled in (a).

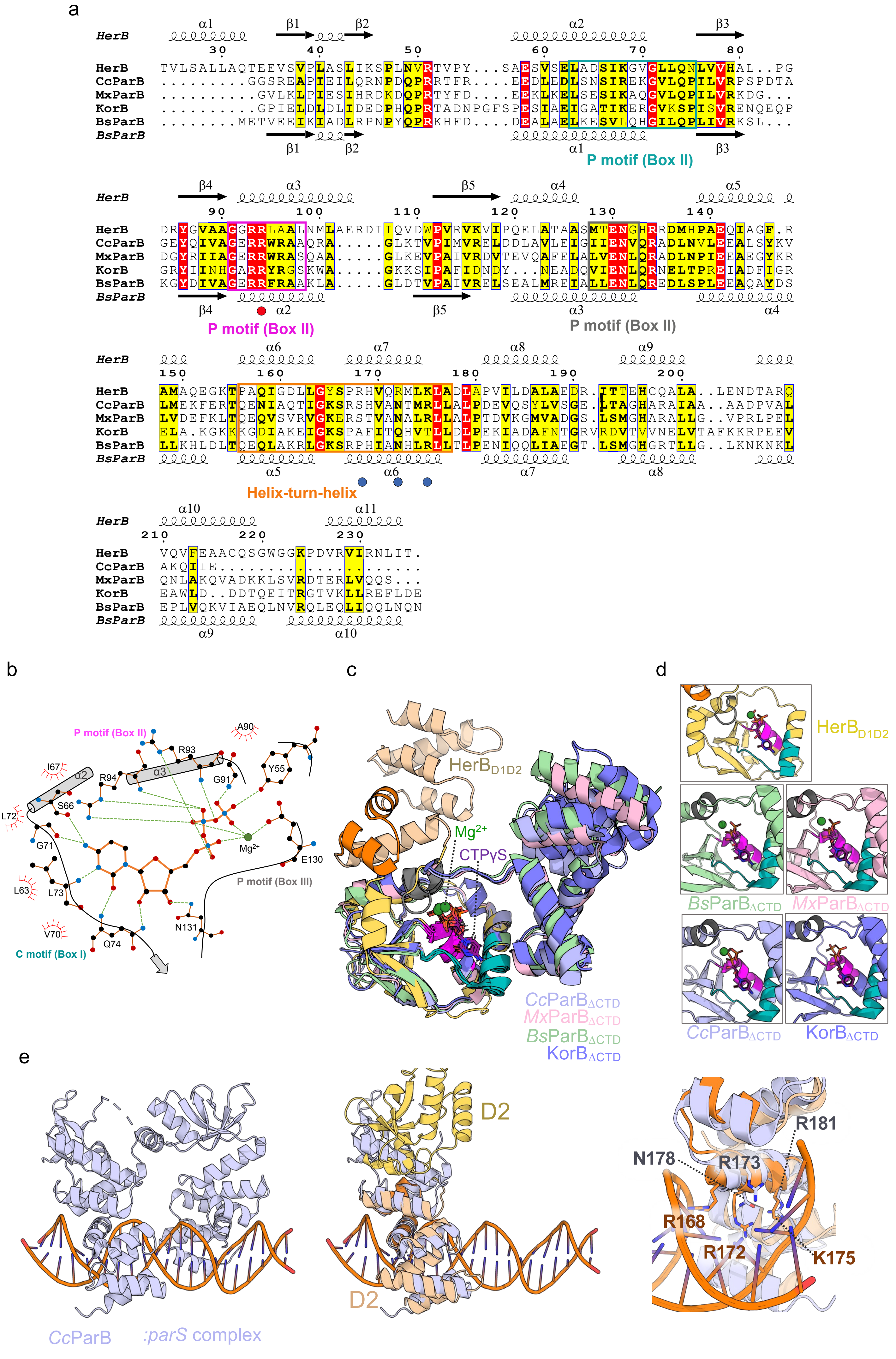

Extended Data Fig. 5

**Extended Data Figure 5. Structural analysis of HerB$_{D1D2}$-CTPγS**

(**a**) Structure-based sequence alignment generated with Fold-Mason of HerB$_{D1D2}$ and ParB sequences from *Caulobacter crescentus* (*Cc*ParB, pdb 7BM8), *Myxococcus xanthus* (*Mc*ParB, pdb 7BNK), *Bacillus subtilis* (*Bs*ParB, pdb 6SDK) and KorB from *Escherichia coli* (KorB, pdb 8QA8). Secondary structure elements of HerB$_{D1D2}$ and *Bs*ParB are indicated above and below the alignment, respectively. Box I, II and III motifs of CTP binding identified in ParBs are indicated by boxes coloured in teal, magenta and grey, respectively. The helix-turn-helix is indicated by an orange box. Residues mutated in the study are indicated by a dot below the alignment: red for CTP binding and blue for putative DNA binding residues. (**b**) protein-ligand interaction map of CTPγS with HerB$_{D1D2}$. Hydrogen bonds are shown as dashed green lines, hydrophobic interactions as red semi-circles and the $Mg^{2+}$ ion in dark green. BoxI, II and III motifs of CTP binding identified in ParBs are indicated (**c**) Structural superimposition of HerBD1D2-CTPγS with the monomer of ParBΔCTD:CTPγS complex from C. crescentus (PDB:7BM8) colored in pale blue, the monomer of ParBΔCTD:CTPγS of *Myoxococcus xanthus* (PDB:7BNR) in pale pink, the monomer of ParBΔCTD:CTPγS of *Bacillus subtilis* (PDB:6SDK) in pale green, and the monomer of KorBΔCTD:CTPγS of RK2 plasmid of *Escherichia coli* (PDB:8QA8) in blue slate. (**d**) Individual zoom of the binding site for each structure of (c). (**e**) Left panel: ribbon representation of *Cc*ParB complex with ParS pdb 6T1F; Jalal *et al.*, 2021) used to superimpose HerB$_{D1D2}$ structure via its HTH domain (middle panel). Right Panel: identification of R168, R172 and K175 from HerB putatively involved in DNA binding. Residues side chains are represented as ball and sticks together with their corresponding residues in *Cc*ParB.

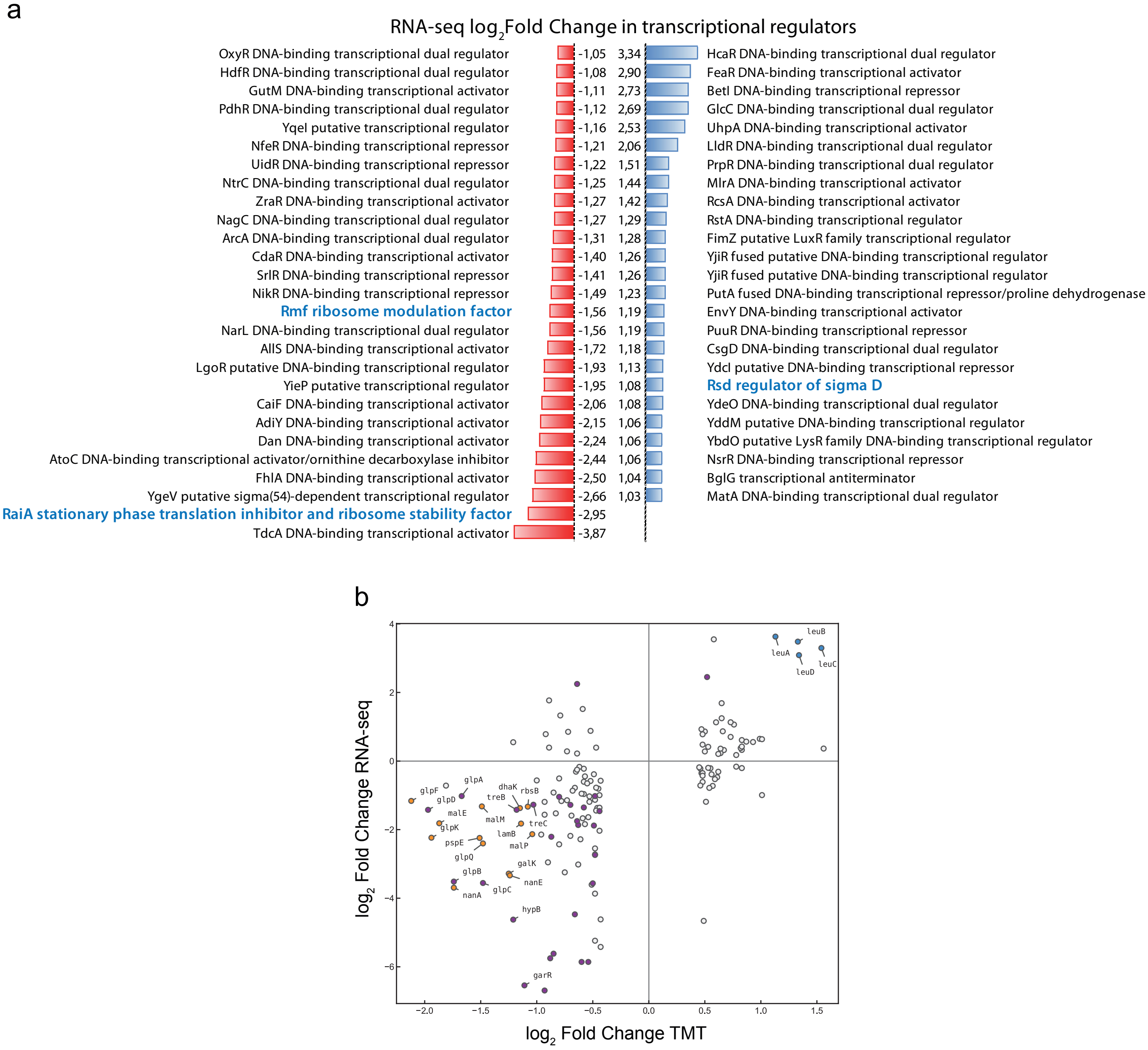

Extended Data Fig. 6

**Extended Data Figure 6.**

(**a**) RNA-seq data focused on transcriptional regulators which expressions are downregulated (red) or upregulated (blue) in the presence of HerB. Three additional genes, *rmf*, *raiA* and *rsd* are shown in blue and discussed in the main text. (**b**) Correlation plot between RNA-seq and TMT data revealing the correlation between genes that are up- or downregulated and proteins that are over- or underrepresented in the presence of HerB.

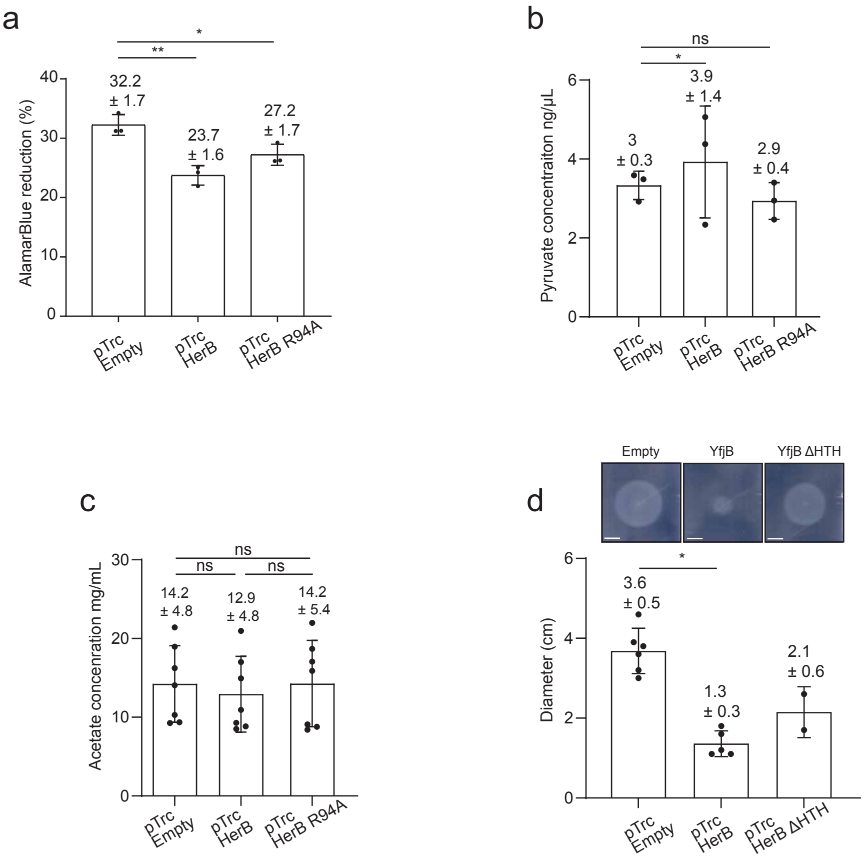

Extended Data Fig. 7

**Extended Data Figure 7.**

(**a**) Histogram showing the AlamarBlue reduction % of strains depending if they are plasmid free or harbouring an empty pTrc99a plasmid, a pTrc99a *herB* plasmid or a double mutant. Mean and SD are calculated based on three individual experiments (black dots). (**b**) Histogram showing the pyruvate production by strains depending if they are plasmid free or harbouring an empty pTrc99a plasmid, a pTrc99a *herB* plasmid, a pTrc99a *herB* plasmid deleted of the HTH domain or harbouring a single mutation on the CTP motif. Mean and SD are calculated based on at least three individual experiments (black dots). (**c**) Histogram showing the acetate production by strains depending if they are plasmid free or harbouring an empty pTrc99a plasmid, a pTrc99a *herB* plasmid, a pTrc99a *herB* plasmid deleted of the HTH domain or harbouring a single mutation on the CTP motif. Mean and SD are calculated based on seven individual experiments (black dots). (**d**) Histogram showing the diameter of strains grown on M9Casa Agar 0.2% medium and depending if they are plasmid free or harbouring an empty pTrc99a plasmid, a pTrc99a *herB* plasmid or a pTrc99a *herB* plasmid deleted of the HTH domain. Images examples are shown on top. Mean and SD are calculated based on at least three individual experiments (black dots). Scale bar 1cm.

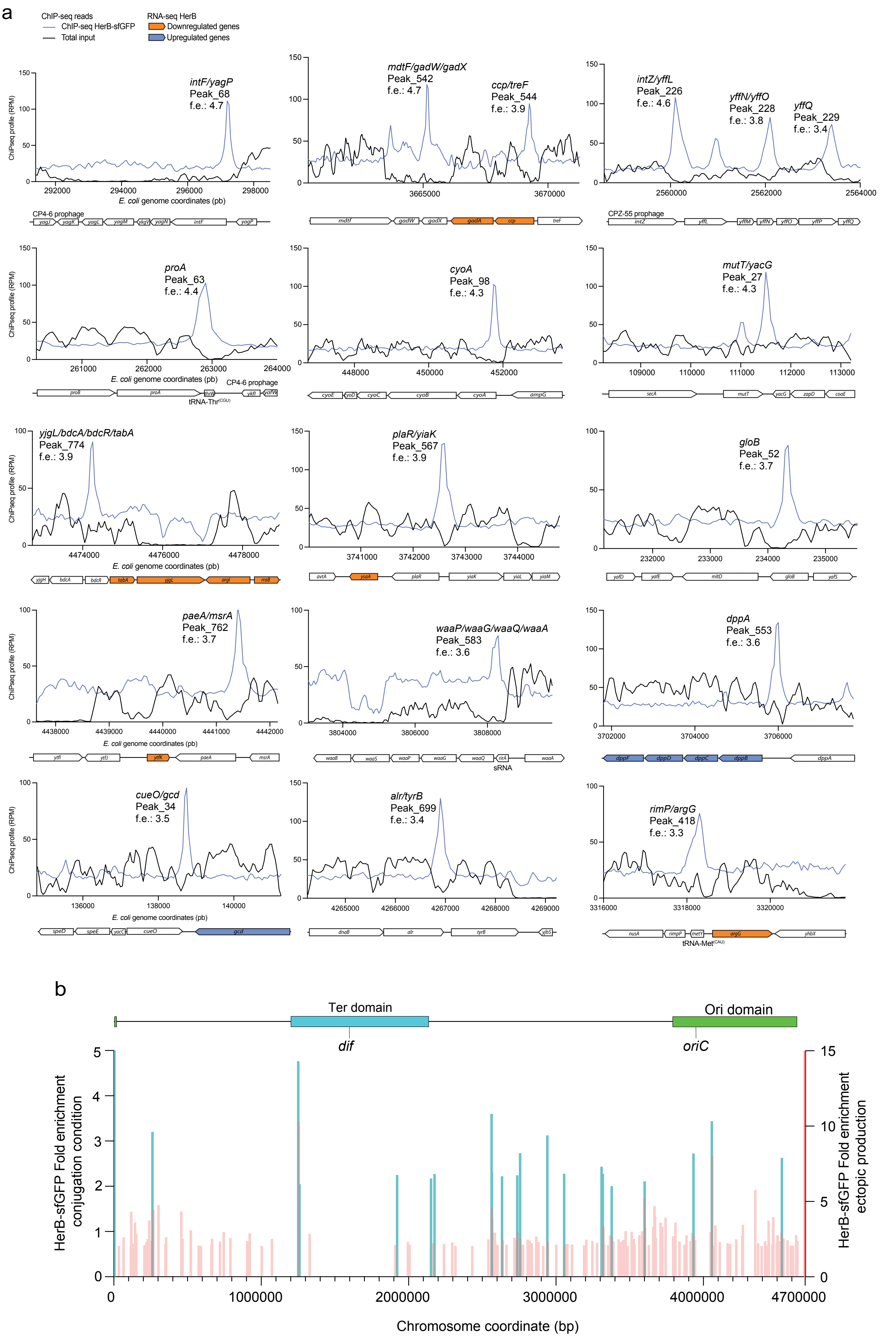

Extended Data Fig. 8

**Extended Data Figure 8.**

(**a**) Genome-browser views showing 15 of the most enriched HerB ChIP-seq peaks. Read density is represented as reads per million (rpm). The names of the genes and the peak number associated with each peak and the corresponding fold enrichment (f.e.) are indicated. Schematics of the corresponding genomic regions are shown below. Genes significantly downregulated or upregulated in the RNA-seq dataset are coloured orange and blue, respectively. (**b**) Distribution and fold enrichment (f.e.) of HerB ChIP-seq peaks (total input-subtracted) in trabnsconjugants having acquired the F *herB* plasmid mapped onto the linearized *E. coli* chromosome. The boundaries of the Ori and Ter macrodomains are indicated above. Merge with the ChIP-seq peaks during ectopic HerB production (corresponding to Fig. 3c) is shown.

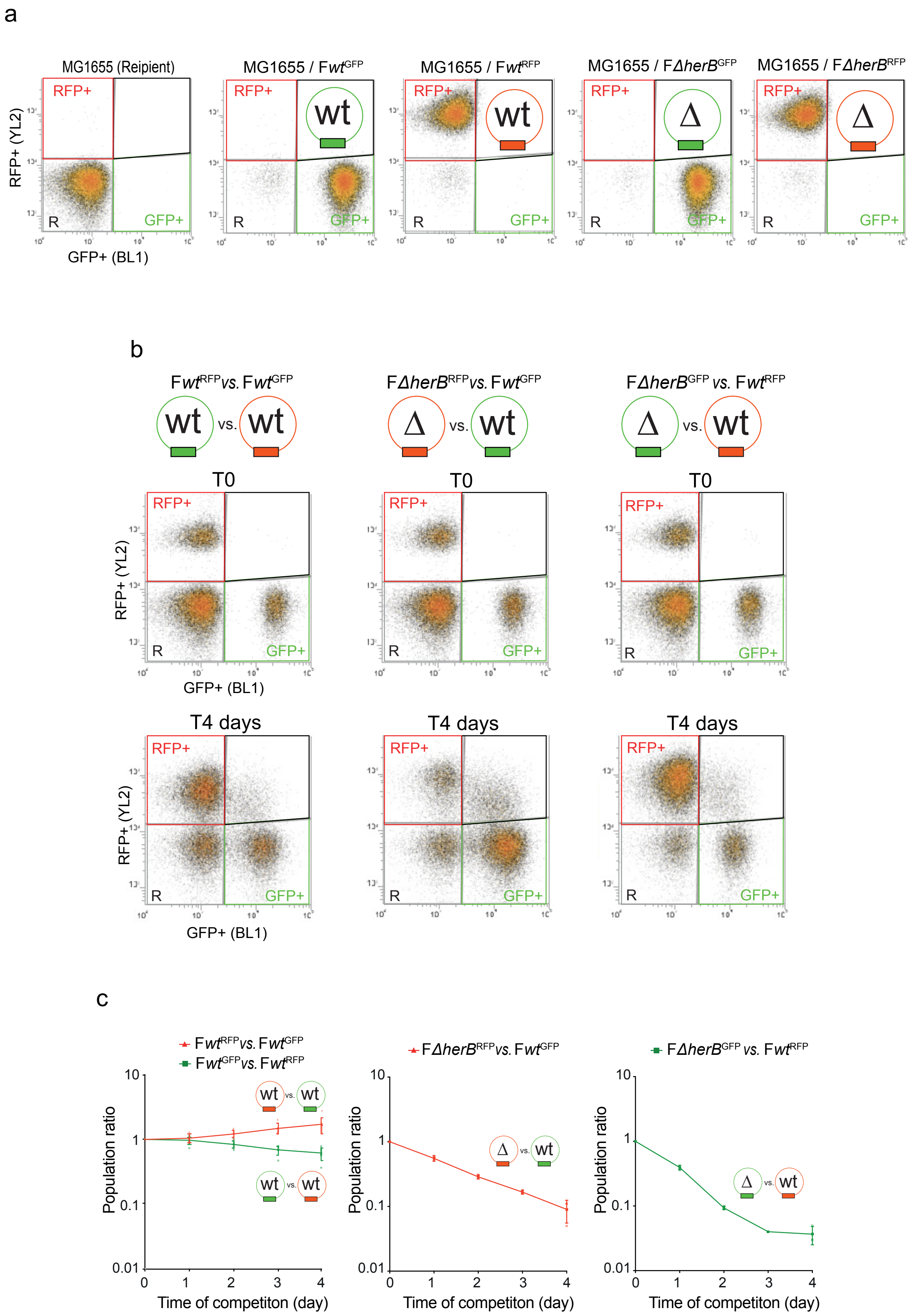

Extended Data Fig. 9

**Extended Data Figure 9. Conjugation-competition assay using flow cytometry. (a)** Flow cytometry analysis of pure populations of F*wt*$^{\mathrm{RFP}}$**,** F*wt*$^{\mathrm{GFP}}$**,** F*ΔherB*$^{\mathrm{RFP,}}$ F*ΔherB*$^{\mathrm{GFP}}$ plasmid donor cells, together with unlabelled recipient cells. Red fluorescence corresponds to RFP, mScarlet2, and green fluorescence corresponds to GFP, mChartreuse. The corresponding analysis gates used for population quantification are shown as grey, red, and green boxes. A total of 100,000 cells were analysed for each sample. **(b)** Flow cytometry profiles of conjugation mixtures F*wt*$^{\mathrm{RFP}}$ *versus* F*wt*$^{\mathrm{GFP}}$; F*ΔherB*$^{\mathrm{RFP}}$ *versus* F*wt*$^{\mathrm{GFP}}$ and F*ΔherB*$^{\mathrm{GFP}}$ *versus* F*wt*$^{\mathrm{RFP}}$, showing the three detected populations, including non-fluorescent recipient cells, RFP-positive cells, and GFP-positive cells. Profiles are shown at T0 and after 2 and 4 days of conjugation competition. A total of 100,000 cells were analysed for each sample. **(c)** Ratios of RFP-positive to GFP-positive cells measured daily over the 4-day conjugation-competition experiment. In the F*wt*$^{\mathrm{RFP}}$ *versus* F*wt*$^{\mathrm{GFP}}$ competition, the GFP marker, mChartreuse, was associated with a minor but detectable disadvantage compared with the RFP marker, mScarlet2. In Fig. 4f, the F*ΔherB*$^{\mathrm{RFP}}$ *versus* F*wt*$^{\mathrm{GFP}}$ and F*ΔherB*$^{\mathrm{GFP}}$ *versus* F*wt*$^{\mathrm{RFP}}$ results were normalized to the F*wt*$^{\mathrm{RFP}}$ *versus* F*wt*$^{\mathrm{GFP}}$ data to compensate for this marker-dependent effect.

| Data collection | YfjB |
|---|---|
| Voltage (kV) | 300 |
| Camera | Gatan K3 |
| Magnification | 105 000 |
| Nominal defocus range (µm) | -0.8 - -2.2 |
| Exposure time (s) | 2.55 |
| Electron exposure (e-/Å$^2$) | 49.87 |
| Number of frames collected | 50 |
| Pixel size (Å) | 0.84 |
| Micrographs (no.) | 8312 |
| Initial particle (no) | 7 306,137 |
| Total particle selected (no.) | 186,180 |
| **Refinement** | |
| Map resolution (Å), 0.143 FSC | 3.5 |
| Map resolution (Å), 0.5 FSC | 4.1 |
| Map sharpening *B* factor (Å) | 202.6 |
| **Model composition** | |
| Non-hydrogen atoms | 3254 |
| Protein residues | 424 |
| Ligands | 0 |
| ***B* factors (Å$^2$)** | |
| Proteins | 76.85 |
| *R.m.s* deviations | |
| Bond lengths (Å) | 0.004 |
| Bond angles (°) | 0.674 |
| **Validation** | |
| MolProbity score | 1.54 |
| Clash score | 10.62 |
| Rotamers outliers (%) | 0.29 |
| Cb outliers (%) | 0 |
| **Ramachandran plot** | |
| Favored (%) | 98.1 |
| Allowed (%) | 1.9 |
| Disallowed (%) | 0 |
| **Model versus map** | |
| Correlation coefficient (mask) | 0.69 |
| Correlation coefficient (box) | 0.64 |
| Correlation coefficient (peaks) | 0.52 |
| Correlation coefficient (volume) | 0.68 |

**Supplementary Table 1. Cryo-EM data collection, refinement, and validation statistic**

| | $HerB_{D1D2}$:CTPγ-S |
|---|---|
| **Resolution range** | 32.34 - 2.58 (2.95 - 2.58) |
| **Space group** | P 1 21 1 |
| **Unit cell** | 32.383 88.832 45.805 90 92.78 90 |
| **Total reflections** | 56881 (19810) |
| **Unique reflections** | 8117 (2693) |
| **Multiplicity** | 7.0 (7.4) |
| **Completeness (%)** | 98.83 (98.50) |
| **Mean I/sigma(I)** | 10.07 (2.07) |
| **Wilson B-factor** | 57.50 |
| **R-merge** | 0.12 (0.95) |
| **R-meas** | 0.13 (1.02) |
| **R-pim** | 0.05 (0.37) |
| **CC1/2** | 0.99 (0.80) |
| **CC*** | 0.99 (0.94) |
| **Reflections used in refinement** | 8113 (2694) |
| **Reflections used for R-free** | 384 (140) |
| **R-work** | 0.196 (0.269) |
| **R-free** | 0.223 (0.323) |
| **Number of non-hydrogen atoms** | 1638 |
| **macromolecules** | 1605 |
| **ligands** | 30 |
| **solvent** | 3 |
| **Protein residues** | 212 |
| **RMS(bonds)** | 0.004 |
| **RMS(angles)** | 0.66 |
| **Ramachandran favored (%)** | 100.00 |
| **Ramachandran allowed (%)** | 0.00 |
| **Ramachandran outliers (%)** | 0.00 |
| **Rotamer outliers (%)** | 0.00 |
| **Clashscore** | 3.04 |
| **Average B-factor** | 62.64 |
| **macromolecules** | 62.64 |
| **ligands** | 63.28 |
| **solvent** | 54.61 |

Statistics for the highest-resolution shell are shown in parentheses.

**Supplementary Table 2. Data collection and refinement statistics X-ray crystallography.**

**Supplementary Table 3. RNA-seq data, ChIP-seq data and correlation between the two.**
This table was submitted independently as an excel file.

| Strain | Relevant genotype[a, b] | Source or reference[a] |
|---|---|---|
| **MG1655 and derivatives** | | |
| **MG1655** | E. coli K-12 $\lambda^-$, *rph-1* | Coli Genetic Stock Center (CGSC) #6300 |
| **MS388** | MG1655 *rpsL* ($St^R$) (*phi80+)* | Gift from F. Cornet |
| **MGT** | MG1655 *rpsL* | Gift from F. Cornet |
| **TB28** | MG1655 *ΔlacIZYA* | Bernhardt and de Boer, 2004 |
| **LY5** | MS388 / F-Tn*10* | Conjugation K603 x MS388 |
| **LY110** | MS388 / F-Tn*10-parS*$_{PMT1}$*-FRT-cat-FRT* | λred *parS*$_{PMT1}$*-FRT-cat-FRT* insertion at the F plasmid intergenic *ygeB-ygfA locus* (OL1/OL2) |
| **LY945** | MS388 *ilvA::cat* | MS388 x P1.LY929 |
| **LY112** | MS388 / F-Tn*10-parS*$_{PMT1}$*-FRT-cat-FRT* | Conjugation LY110 × MS388 to $St^R$ $Tc^R$ |
| **LY118** | MS388 *ssb-ypet-FRT-kan-FRT* | MS388 × P1.RRL505 to $Kn^R$ |
| **LY119** | MS388 *hupA-mCherry-FRT-kan-FRT* | MS388 x P1.Ox468 to $Kn^R$ |
| **LY124** | MS388 / F-Tn*10- parS*$_{PMT1}$*-FRT* | Derivative of LY112, *kan* removed via pCP20 |
| **LY128** | MS388 *ssb-ypet-FRT* | Derivative of LY117, *kan* removed via pCP20 |
| **LY248** | MS388 *hupA-mCherry-FRT* | Derivative of LY119, *kan* removed via pCP20 |
| **LY318** | MS388 / pSN70 | pSN70 × MS388 to $Ap^R$ |
| **LY358** | MS388 *ssb-ypet-FRT* / pSN70 | pSN70 × LY128 to $Ap^R$ |
| **LY643** | TB28 *ilvA::ISceICS-FRT-codA::ISceICS-FRT-yeiU::ISceICS-FRT-ydeO::ISceICS-FRT* | TB28 with 4 cutting site recognised by ISceI |
| **LY735** | MS388 / F-Tn*10-parS*$_{PMT1}$*-FRT-herB-sfGFP-FRT-kan-FRT* | Conjugation LY731 x MS388 to $St^R$ $Tc^R$ |
| **LY792** | MS388 / F-Tn*10-parS*$_{PMT1}$*-FRT-herB-sfGFP-FRT* | Derivative of LY735, *kan* removed via pCP20 |
| **LY1007** | MS388 *ssb-ypet-FRT /* F-Tn*10-parS*$_{PMT1}$*-FRT* | Conjugation LY162 x LY128 $St^R$ $Tc^R$ |
| **LY1266** | MS388 / F-*Tn10-parS*$_{PMT1}$*-FRT, ΔherB::FRT-kan-FRT* | Conjugation LY1242 x MS388 to $St^R$ $Tc^R$ $Kn^R$ |
| **LY1280** | MS388 / F-*Tn10-parS*$_{PMT1}$*-FRT-ΔherB-FRT* | Derivative of LY1266, *kan* removed via pCP20 |

| | | |
|---|---|---|
| **LY1289** | MS388 F-*Tn10-ParS$_{PMT1}$-herB-mcherry-FRT-kn-FRT* | Conjugation LY1278 x MS388 to $Kn^R$ |
| **LY1297** | MG1655 *hupA-GFP-FRT-kan-FRT* | Lab collection |
| **LY1301** | MS388 / F- *Tn10-ParSPMT1-FRT-herB-mcherry-FRT* | Derivative of LY1289, *kan* removed via pCP20 |
| **LY1343** | MS388 / pBG39 | pBG39 x MS388 to $Ap^R$ |
| **LY1344** | MS388 / F-Tn*10* / pBG39 | pBG39 x LY5 to $Ap^R$ |
| **LY1416** | MS388 / pTrc99a | pTrc99a x MS388 to $Ap^R$ |
| **LY1468** | MS388 *hupA-mCherry-FRT* / pBG39 | pBG39 x LY248 to $Ap^R$ |
| **LY1476** | MS388 / pBG49 | pBG49 x MS388 to $Ap^R$ |
| **LY1513** | MS388 *ssb-ypet-kan* / F-Tn*10-parS$_{PMT1}$-FRT-ΔherB-FRT* | LY1280 x P1.LY118 to $Kn^R$ |
| **LY1871** | MS388 / pCV1 | pCV1 x MS388 to $Ap^R$ |
| **LY1936** | MS388 / pCV6 | pCV6 x MS388 to $Ap^R$ |
| **LY1937** | MS388 / pCV7 | pCV7 x MS388 to $Ap^R$ |
| **LY1938** | MS388 / pCV8 | pCV8 x MS388 to $Ap^R$ |
| **LY2107** | MS388 / pCV18 | pCV18 x MS388 to $Ap^R$ |
| **LY2125** | MS388 / pCV20 | pCV20 x MS388 to $Ap^R$ |
| **LY2209** | MS388 / pCV24 | pCV24 x MS388 to $Ap^R$ |
| **LY2219** | MS388 *hupA-mCherry-FRT* / pTrc99a-*sfGFP* | pTrc99a-*sfGFP* x LY248 to $Ap^R$ |
| **LY3212** | DLT4151 | Gift from J. Rech, Centre de Biologie Intégrative. Derivative of LY643 to add Pcp18::*araE-FRT-kan-FRT-recA-* |
| **LY3305** | MGT *ilvA::cat* | MGT x P1.LY945 to $Cm^R$ |
| **LY3372** | MG1655 *rpsL* ($St^R$) | Spontaneous mutation of K43R (AAA to AGA) on MGT to $St^R$ |
| **LY3399** | MGT St / F-*Tn10-ParS$_{PMT1}$-FRT* | Conjugation LY162 x LY3372 to $St^R$ $Tc^R$ |
| **LY3450** | MGT St / pTrc99a-herB | pTrc99a-*herB* x MGT St to $Ap^R$ |
| **LY3473** | DLT4151 | Derivative of LY3213, *kan* removed via pCP20 |
| **LY3601** | DLT4151-*HupA-mCherry-FRT-kan-FRT* | P1 LY119 x LY3473 to $Kn^R$ |
| **LY3728** | MGT *ilvA::cat* / F-*Tn10-parS$_{PMT1}$-FRT* | Conjugation LY162 x LY3305 to $St^R$ $Tc^R$ |
| **LY3984** | MGT *ilvA::cat* / F-*Tn10-parS$_{PMT1}$-FT-ΔherB-FRT* | Conjugation LY1280 x LY3305 to $Tc^R$ |

| | | |
|---|---|---|
| **LY3988** | MGT *ilvA::cat* / pTrc99a | pTrc99a x LY3305 to 3305 to $Ap^R$ |
| **LY3989** | MGT *ilvA::cat* / pTrc99a-*herB* | pTrc99a-herB x LY3305 to $Ap^R$ |
| **LY3990** | MGT St / FTn10-parS$_{PMT1}$-FRT-Δ*herB*-FRT | Conjugation LY3984 x LY3372 to $St^R$ $Tc^R$ |
| **LY4412** | DLT4151-*HupA-mCherry-FRT-kan-FRT* / pSCe / pTrc99a-*herB-sfgfp* | pTrc99a-*herB-sfgfp* x LY4711 to $Ap^R$ |
| **LY4463** | MS388 *hupA-mCherry-FRT* / pTrc99a-*herB-R168A-R172A-sfGFP* | pTrc99a-*herB-R168A-R172A-sfGFP* x LY248 to $Ap^R$ |
| **LY4464** | MS388 *hupA-mCherry-FRT* / pTrc99a- *herB-R168A-R172A-K175A-sfGFP* | pTrc99a-*herB-R168A-R172A-K175A-sfGFP* x LY248 to $Ap^R$ |
| **LY4508** | MS388 *hupA-mCherry-FRT* / pTrc99a- *herB-E130A-sfGFP* | pTrc99a-*herB-E130A-sfGFP* x LY248 to $Ap^R$ |
| **LY4669** | MGT St / F-*Tn10 repE::Pbiofab-mchartreuse-FRT* | λred *Pbiofab-mchartreuse* fusion at endogenous F plasmid locus (OL117/817) in DY330. Conjugation to MGT St to $Tc^R$. *kan* removed via pCP20. |
| **LY4711** | DLT4151-*HupA-FRT-kan-FRT* / pSCe | pSCLe x LY3601 to $Cm^R$ |
| **LY4727** | MGT St / F-*Tn10 repE::Pbiofab-mscarletI3-FRT-kan-FRT* | Conjugation LY4716 x MGT St to $Tc^R$ |
| **LY4747** | MGT St / F-*Tn10 repE::Pbiofab-mscarletI3-FRT* | Derivative of LY4727, *kan* removed via pCP20 |
| **LY5184** | F-*Tn10-parS$_{PMT1}$-FRT-ΔherB* | 2 steps recombination using SacB suicide plasmid pKanSac-F-ΔherB. |
| **LY5294** | MGT St / F-*Tn10 repE::Pbiofab-mchartreuse-FRT-kan-FRT-parS$_{PMT1}$-FRT-ΔherB* | Conjugation LY5292 x MGT St to $Tc^R$ |
| **LY5295** | MGT St / F-*Tn10 repE::Pbiofab-mscarletI3-FRT-kan-FRT-parS$_{PMT1}$-FRT-ΔherB* | Conjugation LY5293 x MGT St to $Tc^R$ |
| **LY5296** | MGT St / F-*Tn10 repE::Pbiofab-mchartreuse-FRT-parS$_{PMT1}$-FRT-ΔherB* | Derivative of LY5294, *kan* removed via pCP20 |
| **LY5297** | MGT St / F-*Tn10 repE::Pbiofab-mscarletI3-FRT-parS$_{PMT1}$-FRT-ΔherB* | Derivative of LY5295, *kan* removed via pCP20 |
| **Other genetic backgrounds** | | |
| **BL21 (DE3)** | $F^-$*omp*T *hsd*$S_B$ ($r_B^-$, $m_B^-$) *GAldcmrne*131 (DE3) | Thermo fisher (#C606010) |

| | | |
|---|---|---|
| **Ox468** | W1485 *hupA-mCherry-FRT-kan-FRT* | Gift from F. Cornet |
| **K603** | F+ [F1-10(Tn10)], *thr-1, araC14, leuB6(Am), lacY1, glnX44(AS), galK2(Oc), galT22, λ-, ΔtrpE63, xylA5, mtl-1, thiE1,* | Coli Genetic Stock Center (CGSC) #6451 |
| **DH5α** | F– *endA1 glnV44 thi-1 recA1 relA1 gyrA96 deoR nupG purB20* φ80d*lacZ*ΔM15 Δ(*lacZYA-argF*)U169, hsdR17(*rK–mK+*), λ | Lab collection |
| **DY330** | W3110 Δ*lacU169, gal490, λcI857,* Δ*(cro-bioA)* | Yu *et al.*, 2000 |
| **RRL505** | AB1157 Δ*acrA::FRT-ssb-ypet-FRT-kan-FRT* | Gift from R. Reyes-Lamothe, McGill University |
| **LY162** | DY330 / F-Tn*10-parS$_{PMT1}$-FRT* | Conjugation LY124 × DY330 to $Tc^R$ |
| **LY156** | DY330 / F- Tn*10* | Conjugation LY5 x DY330 to $Tc^R$ |
| **LY731** | DY330 / F- Tn*10*-ParS$_{PMT1}$-FRT-*herB-sfGFP-FRT-kan-FRT* | λred *herB-sfgfp* fusion at the endogenous F plasmid locus (OL418/OL607) |
| **LY929** | DY330 *ilvA::cat* | λred *ilvA::cat* at chromosome locus (OL418/OL419) |
| **LY1242** | DY330 / F-Tn*10-parS$_{PMT1}$-FRT-ΔherB::FRT-kan-FRT* | λred Δ*herB::kan* construct at the endogenous F plasmid *locus* (OL606/OL607) |
| **LY1333** | DH5α / pBG39 | Gibson assembly *herB-sfGFP* construct from LY792 on pTrc99a plasmid (OL604/OL605 and OL602/ OL603) |
| **LY1279** | DY330 / F-Tn*10-ParS$_{PMT1}$-FRT-herB-mcherry-FRT-kan-FRT* | λred *herB-mCherry* fusion at the endogenous F plasmid locus (OL418/OL607) |
| **LY2035** | BL21 DE3 / pCV15 | pCV15 x BL21 star to $Ap^R$ |
| **LY4716** | DY330 / F-*Tn10 repE::P$_{biofab}$-mscarletI3-FRT-kan-FRT* | λred *Pbiofab-mscarletI3* fusion at endogenous F plasmid locus (OL117/688) |
| **LY5292** | DY330 / F-*Tn10 repE::P$_{biofab}$-mchartreuse-FRT-kan-FRT-parS$_{PMT1}$-FRT-*Δ*herB* | λred *Pbiofab-mchartreuse* fusion at endogenous F plasmid locus (OL117/817) |
| **LY5293** | DY330 / F-*Tn10 repE::P$_{biofab}$-mscarletI3-FRT-kan-FRT-parS$_{PMT1}$-FRT-*Δ*herB* | λred *Pbiofab-mscarletI3* fusion at endogenous F plasmid locus (OL117/688) |
| **LY5308** | BL21 DE3/ pET151-*herB$_{D1D2}$* | pET151D-*herB$_{D1D2}$* x BL21 star to $Ap^R$ |

[a] The abbreviation *kan* refer to insertions conferring resistance to kanamycin ($Kn^R$) and the abbreviation *cat* to chloramphenicol ($Cm^R$). $Ap^R$ refers to ampicillin resistance. $St^R$ refers to streptomycin resistance. $Tc^R$ refers to tetracycline resistance. *FRT* refers to the FLP site-specific recombination site.
[b]Tn*10* transposon is located in the intergenic region *ybdB-ybfA* on the F plasmid.
[c] *sfgfp* gene encodes the superfolder Green Fluorescent Protein sfGFP

**Supplementary Table 4. Strains list**

| Name | **Construct and Usage**[a] | **Source or reference** |
|---|---|---|
| pCP20 | Flp expression plasmid, $Ap^R$, $Cm^R$, ts | Datsenko *et al.*, 2000 |
| p*mcherry-parB*$_{PMT1}$ (pSN70) | IPTG inducible expression of N-terminal fusion *mcherry-ParB*$_{PMT1}$ | Nolivos *et al.*, 2019 |
| pR6K-sfGFP | Carries *sfGFP-FRT-kan-FRT* used for several C-terminal fusion by λred | Nolivos *et al.*, 2019 |
| pR6K-$P_{biofab}$-*mchartreuse* | Carries *Pbiofab-mchartreuse-FRT-kan-FRT* used for insertion by λred | |
| pR6K-$P_{biofa}$*b-mscarletI3* | Carries *Pbiofab--mscarletI3-FRT-kan-FRT* used for insertion by λred | |
| pROD62 | Carries *FRT-kan-FRT* used for deletion by λred | Gift from R. Reyes-Lamothe, McGill University |
| pSEVA321 | Carries *FRT-cat-FRT* used for deletion by λred | Gift from V. de Lorenzo, Centro Nacional de Biotecnología |
| pTrc99a | Lab construct (GE HEALTH CARE) | |
| pTrc99a-*herB-sfgfp* (pBG39) | Gibson assembly *herB-sfGFP* construct from LY792 on pTrc99a plasmid (OL604/OL605) and (OL602/ OL603) | |
| pTrc99a-*herB-mcherry* (pBG49) | Gibson assembly *herB-mCherry* construct from LY1301 on pTrc99a plasmid (OL604/OL653) and OL602/ OL603) | |
| pTrc99a-*herB* | *sfGFP* deletion by PCR-ligation of pBG39 (OL602/OL776) | |
| pTrc99a-*herB*$_{D1D2}$-*sfgfp* | Deletion by PCR-ligation of pBG39 (OL922/OL1685) | |
| pTrc99a-*herB*$_{D1D2D3}$- *sfgfp* | Deletion by PCR-ligation of pBG39 (OL922/OL1686) | |
| pTrc99a-*herB*$_{D3D4}$- *sfgfp* | Deletion by PCR-ligation of pBG39 (OL1688/OL675) | |
| pTrc99a-h*erB*$_{D4}$-*sf sfgfp* | Deletion by PCR-ligation of pBG39 (OL1687/OL675) | |
| pTrc99a-h*erB-ΔHTH* (pCV1) | Phosphorylation-ligation of pTrc99a-*herB* (OL903/OL904) | |

| | | |
|---|---|---|
| pTrc99a-*herB-R168A-sfgfp* (pCV6) | Directed mutagenesis (OL906/OL907) | |
| pTrc99a-*herB-R177A-sfgfp* (pCV7) | Directed mutagenesis (OL908/OL909) | |
| pTrc99a-h*erB-K175A- sfgfp* (pCV8) | Directed mutagenesis (OL910/OL911) | |
| pTrc99a-*herB-R168-R172A-sfgfp* | Directed mutagenesis (OL912/OL907) | |
| pTrc99a-*herB-R168-R172A-K175A- sfgfp* | Directed mutagenesis (OL3185/OL907) | |
| pTrc99a *herB-R94A-sfgfp* (pCV18) | Directed mutagenesis (OL1006/OL1007) | |
| pET151D-*herB* (pCV15) | PCR of *herB* (OL932/OL933) from pTrc99a-*herB* and mix Topo151 kit | |
| pTrc99a-*herB-R94A-ΔHTH* (pCV20) | Directed mutagenesis (OL1006/OL1007) on pCV1 | |
| pTrc99a-*herB-R94A* (pCV24) | Directed mutagenesis (OL1006/OL1007) | |
| pTrc99a-*herB-E130A-sfgfp* | Directed mutagenesis (OL2549/OL3181) | |
| pTrc99a-*sfgfp* | *herB* deletion by PCR-ligation of pBG39 (OL675/OL775) | |
| pET151D-$herB_{D1D2}$ | PCR of pCV15 (OL3163/OL3168) and mix TOPO151 kit | |
| pKanSac | Lab construct. SacB suicide plasmid with non-replicative R6K with SacB and LacZα, $Kn^R$. | |
| pKanSac-FΔ*herB* | For 2 steps recombination with SacB suicide plasmid for ***herB*** deletion. Gibson assembly construct from LY3399 (OL2971/2972) and (2973/2974) on pKanSaC plasmid (OL2301/2302). | |
| **pSCe** | Arabinose inducible expression of I-SceI endonuclease | Fraikin and Van Melderen, 2024 |
| **F-Tn*10* conjugative plasmid (from K603) derivatives** | | |
| F*wt* | F-Tn*10* with $parS_{PMT1}$ inserted at the intergenic *ygeB-ygfA locus* | |

| | |
|---|---|
| F *herB-sfGFP* | F-Tn*10* $parS_{PMT1}$ with *herB-sfgfp* translational fusion at the endogenous locus |
| F *herB-mCherry* | F-Tn*10* $parS_{PMT1}$ with *herB-sfgfp* translational fusion at the endogenous locus |
| F Δ*herB* | F-Tn*10* $parS_{PMT1}$ with *herB* deletion |
| F *repE::*$P_{biofa}$*b-mchartreuse* | F-Tn*10* $parS_{PMT1}$ wih insertion of *Pbiofab-mchartreuse* in the *repE-sopA* intergenic region |
| F *repE::*$P_{biofab}$*-mscarletI3* | F-Tn*10* $parS_{PMT1}$ and wih insertion of *Pbiofab-mchartreuse* in the *repE-sopA* intergenic region |
| F *repE::*$P_{biofab}$*-mchartreuse-*Δ*herB* | F-Tn*10* $parS_{PMT1}$ with *herB* deletion and wih insertion of *Pbiofab-mchartreuse* in the *repE-sopA* intergenic region |
| F *repE::*$P_{biofab}$*-mscarletI3-*Δ*herB* | F-Tn*10* $parS_{PMT1}$ with *herB* deletion and wih insertion of *Pbiofab-mscarletI3* in the *repE-sopA* intergenic region |

[a] *sfgfp* gene encodes the superfolder Green Fluorescent Protein sfGFP

**Supplementary Table 5. Plasmids used in this study**

| Name | Sequence | Construct |
|---|---|---|
| **OL1** | GTATTGTCACTCATATGATATTTTTGTGTGTGGC CTTCCAGGTCTGCTATGTGGTGCTATCT | λred *parS<sub></sub>*$_{PMT1}$*-FRT-cat-FRT* insertion at the F plasmid intergenic *ygeB-ygfA locus*. PCR on pGBKD3-parS |
| **OL2** | ACAGGCATTGTCAGATACCGGTTATGCCGCAA AAGCGGCAGATTGTGTAGGCTGGAGCTGC | |
| **OL117** | GAGCATAGCGAGCGAACTGGCGAGGAAGCAAA GAAGAACTCATATGAATATCCTCCTTAG | λred *Pbiofab-mchartreuse-FRT-kan-FRT* insertion or *Pbiofab-mscarletI3* at the endogenous F plasmid locus. PCR on pR6K-*Pbiofab-mchartreuse* plasmid (OL117/688). PCR on pR6K-*Pbiofab-mscarletI3* (OL117/817) |
| **OL418** | GGATGCCACTGAACGTACCGATAACCTGGCTG ATGCCGCCAGCTCGGCTGGCTCCGCTGC | λred *herB-sfgfp* fusion at the endogenous F plasmid *locus*. PCR on pR6K-sfGFP plasmid |
| **OL419** | TGCTGCCGCCCCGTCTCCGGCGGGGCGGTGTGG TTGTTCATATCCTCCTTAGTTCCTATT | |
| **OL495** | GGAAAAATGCCTGATAGCGCTTCGCTTATCAGG CCTACCCGCGTTGATCGGCACGTAAGA | λred *ilvA::cat* at chromosome locus. PCR on PSEVA321 |
| **OL496** | TCTGGAAGATTTTGCCGAACCACAAATGACGTT GTCGCGCGGAAAAGGACAAAAGTCAAA | |
| **OL602** | GATCCTCTAGAGTCGACCTGCAGGC | pTrc99a-*herB-sfgfp* plasmid construct using Gibson assembly at the pTrc99a *locus*. PCR on pTrc99a (OL602/OL603) and PCR on LY792 (OL604/OL605) |
| **OL603** | TCTGGAAGATTTTGCCGAACCACAAATGACGTT GTCGCGCCATATGAATATCCTCCTTAG | |
| **OL604** | AACAATTTCACACAGGAAACAGACCATGTCAG TTACAGAGTCTAA | |
| **OL605** | GCCTGCAGGTCGACTCTAGAGGATCTTATTTGT AGAGCTCATCCA | |
| **OL606** | CGGCAGACAATCATTATTACCGTTTTGGGAGGT ACTAACTGTGTAGGCTGGAGCTGCTTC | λred *kan* insertion at the endogenous F plasmid *herB locus*. PCR on pROD62 plasmid |
| **OL607** | TGCTGCCGCCCCGTCTCCGGCGGGGCGGTGTGG TTGTTCACATATGAATATCCTCCTTAG | |
| **OL653** | GCCTGCAGGTCGACTCTAGAGGATCTTACTTGT ACAGCTCGTCCA | pTrc99a-*herB-mCherry* plasmid construct using Gibson assembly at the pTrc99a *locus*. PCR on pTrc99a (OL602/OL603) and PCR on LY1301 (OL604/OL653) |
| **OL675** | CATGGTCTGTTTCCTGTGTG | *herB* deletion by PCR-ligation of pBG39 (OL675/OL775) |

| | | |
|---|---|---|
| **OL688** | ATTTCTTCTTGCGCTGAGCGTAAGAGCTATCTG ACAGAACCCGACTGGAAGCATCGATAG | λred *Pbiofab-mchartreuse-FRT-kan-FRT* insertion or *Pbiofab-mscarletI3* at the endogenous F plasmid locus. PCR on pR6K-*Pbiofab-mchartreuse* plasmid (OL117/688). |
| **OL775** | TCTAAAGGTGAAGAACTGTTCACC | *herB* deletion by PCR-ligation of pBG39 (OL675/OL775) |
| **OL776** | TCAGGCGGCATCAGCCAGGT | *sfGFP* deletion by PCR-ligation on pBG39 plasmid (OL602/OL776) |
| **OL817** | ATTTCTTCTTGCGCTGAGCGTAAGAGCTATCTG ACAGAACGACAGGTTTCCCGACTGGAA | λred *Pbiofab-mscarletI3-FRT-kan-FRT* at the endogenous F plasmid locus. PCR on pR6K- *Pbiofab-mscarletI3* plasmid (OL117/817). |
| **OL903** | CTGGCAGACCTTGCGCCTGTCATCC | HTH motif deletion (K164 to K175) by PCR-ligation on pTrc99a-*herB* |
| **OL904** | GCCTTCCTGCGCCATTGCACGGA | |
| **OL906** | CTGGGTTATTCGCCCGCCCACGTTCAGCGAATG CTGAAACTGGC | R168A mutation by directed mutagenesis. PCR on pBG39 plasmid |
| **OL907** | GCGGGCGAATAACCCAGCAAATCACCG | |
| **OL908** | CCGCCACGTTCAGGCAATGCTGAAACTGGCAG ACCTTGCGC | R172A mutation by directed mutagenesis. PCR on pBG39 plasmid |
| **OL909** | GCCTGAACGTGGCGGGGCGAATAAC | |
| **OL910** | GTTCAGCGAATGCTGGCACTGGCAGACCTTGCG CCTGT | K175A mutation by directed mutagenesis. PCR on pBG39 plasmid |
| **OL911** | GCCAGCATTCGCTGAACGTGGCGGG | |
| **OL912** | GGGTTATTCGCCCGCCCACGTTCAGGCAATGCT GAAACTGGCAGACCTTGC | R168A-R172A mutation by directed mutagenesis. PCR on pBG39 (OL912/OL907) |
| **OL922** | AGCTCGGCTGGCTCCGCTGC | D3D4 (OL922/OL1685) and D4 (OL922/OL1686) deletion by PCR-ligation on pBG39 |
| **OL932** | CACCATGTCAGTTACAGAGTCTAAGG | pET151D-*herB* plasmid construct with Topo151 kit mix PCR on pTrc99a-*herB* |
| **OL933** | TCAGGCGGCATCAGCCA | |
| **OL1006** | GCCGCAGGTGGTCGCGCACTGGCAGCACTCAA CATGCTG | R94A mutation by directed mutagenesis. PCR on pBG39 plasmid, pTrc99a-*herB* and pCV1 |
| **OL1007** | GCGCGACCACCTGCGGCGACACCGT | |
| **OL1685** | TTCCGTAATCAGATTGCGGATAACCC | $herB_{D3D4}$ deletion by PCR-ligation on pBG39 (OL922/OL1685) |

| | | |
|---|---|---|
| **OL1686** | CTGCACCTGTTCCGTGCGG | *herB*$_{D4}$ deletion by PCR-ligation on pBG39 (OL922/OL1686) |
| **OL1687** | GAAAAAGCGTCAGTGGAGGAAATCAGTC | *herB*$_{D1D2D3}$ deletion by PCR-ligation on pBG39 (OL1687/OL675) |
| **OL1688** | AGTGAAGTGGCGGTGAAGGACAAC | *herB*$_{D1D2}$ deletion by PCR-ligation on pBG39 (OL1688/OL675) |
| **OL2301** | GCAAGATCCGCAGTTCAACC | Suicide plasmid construct by Gibson assembly. PCR on pKanSac (OL2301/2302) and on LY3399 (OL2971/2972) and (OL2973/2974) |
| **OL2302** | TGAATGGCGAATGGCGCTTC | |
| **OL2549** | CAGTCGCCAGCTCCTGCGG | E130A mutation by directed mutagenesis. PCR on pBG39 (OL2549/OL3181) |
| **OL2971** | GAAGCGCCATTCGCCATTCAGTTCGTGGCATTA ACAAGGTC | Suicide plasmid construct by Gibson assembly. PCR on pKanSac (OL2301/2302) and on LY3399 (OL2971/2972) and (OL2973/2974) |
| **OL2972** | CCGGCGGGGCGGTGTGGTTGGTAATAATGATT GTCTGCCGGAC | |
| **OL2973** | GTCCGGCAGACAATCATTATTACCAACCACACC GCCCC | |
| **OL2974** | GGTTGAACTGCGGATCTTGCCCTCAAGCGCCGG G | |
| **OL3163** | CACCATGTCAGTTACAGAGTCTAAGGC | pET151D-*herB*$_{D1D2}$ plasmid construct with Topo151 kit mix PCR on pTrc99a-*herB* |
| **OL3168** | TCAACTTTCCGTAATCAGATTGCGG | |
| **OL3181** | CCGCATCGATGACCGCGAACGGTCATCGTCG | E130A mutation by directed mutagenesis. PCR on pBG39 (OL2549/OL3181) |
| **OL3185** | CTGGGTTATTCGCCCGCCCACGTTCAGGCAATG CTGGCACTGGCAGACCTTGCGCCTG | R168A-R1721-K175A mutation by directed mutagenesis. PCR on pCV7 (OL3185/OL907) |

**Supplementary Table 6. Primers used for strain and plasmid construction**

**Supplementary Video Legends**

**Supplementary Video S1. HerB-sfGFP production and dynamics in transconjugants cells having received the FherB-sfgfp plasmid by conjugation.** Time-lapse microscopy imaging within a microfluidic chamber of a conjugation mix containing F*herB-sfgfp* donors (no fluorescence) and recipient cells producing mCherry-ParB (diffuse red fluorescence). Acquisition of the plasmid in reflected by the formation of a red mCherry-ParB focus in transconjugant cells, followed by the production of green fluorescence HerB-sfGFP, which form bright and dynamic foci. Scale bar 5µm. Time indicated (5 min/frame time intervals over 120 min).

**Supplementary Video S2. Dynamics of nucleoid-associated HerB-sfGFP foci.** Time-lapse microscopy imaging within a microfluidic chamber of cells producing the nucleoid-associated protein HU-mCherry (red fluorescence), and ectopically producing HerB-sfGFP from the pTrc *herBsfgfp* plasmid (green fluorescent foci), growing in M9 CAA glucose medium at 37ºC. Scale bar 1µm. Time indicated (5 min/frame time intervals over 145 min).

**Supplementary Video S3. Dynamics of nucleoid-associated HerB-sfGFP foci upon inhibition of transcription by rifampicin treatment.** Time-lapse microscopy imaging within a microfluidic chamber of cells producing the nucleoid-associated protein HU-mCherry (red fluorescence), and ectopically producing HerB-sfGFP from the pTrc *herBsfgfp* plasmid (green fluorescent foci), growing in M9 CAA glucose medium at 37ºC, before during and after treatment with rifampicin. Scale bar 1µm. Time indicated (5 min/frame time intervals over 300 min).

**Supplementary Video S4. Structural comparison of AlphaFold 3 models of HerB.** The AlphaFold3 models of the HerB dimer were superimposed onto the conserved $HerB_{D4}$ dimer structure. Morphing between each of the five models, as implemented in ChimeraX, was used to illustrate the various conformations adopted by the leg domains ($HerB_{D1D2D3}$). HerB is coloured as in Figure 2a-b: with D1 in yellow, D2 in wheat, D3 in green and D4 in blue.

**Supplementary Video S5. CryoEM structure of $HerB_{D4}$.** The electron density map (contoured at level 0.152) of the $HerB_{D4}$ dimer coloured by chain, with the fitted atomic models of $HerB_{D4}$ chain A (blue) and chain B (grey) reconstructed within the density

**Supplementary Video S6. Structural insight into $HerB_{D1D2}$ molecular functions.** The crystal structure of $HerB_{D1D2}$: CTPγS is represented in cartoon and coloured according to the domains Fig.2a,e). The 2Fo-Fc electron map is shown around the CTPγS (magenta) and $Mg^{2+}$ ion (green sphere) contoured at 1.5σ with the side chain of the residues involved in binding represented by sticks and labelled. The helix-turn-helix motif (HTH) is coloured in orange. A view of the $HerB_{D1D2}$: CTPγS model bound to dsDNA via the HTH is also presented, with the three positively charged residues identified as interacting with the major groove depicted as ball-and-stick models.
